# Rank-Resolved Multi-Engine Docking and Optuna-Optimized Re-Ranking with ProDock for Virtual Screening

**DOI:** 10.64898/2026.07.24.740457

**Authors:** Lai Hoang Son Le, Thanh-An Pham, Ngoc-Tam Nguyen Tran, Phuoc-Chung Van-Nguyen, Tieu-Long Phan, Tuyen Ngoc Truong

## Abstract

False positives in virtual screening often arise when a single docking score or top-ranked pose is treated as sufficient evidence for binding. We extend the previously introduced ProDock software from a database-backed docking platform into a rank-resolved, multi-engine workflow for automated preparation, docking, pose analysis, and optimized re-ranking. The extended workflow combines local docking with GNINA and global docking with DiffDock with pose-level descriptors, namely binding-site occupancy, ligand localization, interaction-fingerprint similarity, and steric clash counts, together with Optuna -based threshold optimization. Across 43 DUDE-Z targets, the archived benchmark outputs reported higher enrichment values for CNN-based GNINA scores after optimization. CNNaffinity PR-AUC changed from 0.197 to 0.294 and LogAUC from 0.708 to 0.763, whereas empirical affinity ROC-AUC changed from 0.770 to 0.758. Structural investigation of re-docked actives showed that re-ranked poses were more native-like, with improved binding-site occupancy, reduced centroid displacement, and greater recovery of co-crystal interactions. The extension provides a reproducible framework for combining complementary docking engines with interpretable pose-level metrics before hit selection, thereby aiding the identification of true-positive candidates in virtual screening.

## 1 Introduction

Molecular docking proposes binding poses between proteins and ligands and scores them to prioritize compounds for experimental testing. Classical engines such as AutoDock Vina ^1,2^ remain widely used because they are fast, interpretable, and scalable. However, a rank-1 pose or single score can be a fragile proxy for binding when active compounds and decoys have similar properties or a binding site admits several plausible interaction modes. Collapsing each ligand to one pose can therefore discard useful structural information, while scores that capture different properties may disagree.

Recent docking workflows increasingly combine physics-inspired search, machine learning scoring, and geometric deep learning. GNINA ^3^ extends Vina -like docking^1,2^ with convolutional neural network scoring and pose assessment, whereas DiffDock ^4^ treats docking as a generative diffusion problem over ligand translations, rotations, and torsions. Other deep-learning docking methods, including EquiBind ^5^ and SurfDock,^6^ further illustrate the rapid movement toward direct pose generation and learned three-dimensional representations. While these methodological developments enable more rigorous virtual screening protocols, they introduce substantial operational overhead. Specifically, macromolecular structures and ligands must be prepared uniformly through manual procedures, different docking programs output disparate scoring metrics, and candidate selection requires sophisticated evaluation rather than simply choosing the first generated pose.

Several platforms have addressed parts of this automation problem. VirtualFlow ^7^ supports ultra-large virtual screens on computing clusters, DockStream ^8^ provides a docking wrapper designed for integration with *de novo* molecular design, and OpenDock ^9^ offers a PyTorch -based docking framework with modular scoring and sampling components. Although these platforms support large-scale docking workflows, they were not primarily designed to analyze multiple pose ranks, and they usually rely on one or a few metrics for virtual-screening candidate selection. We previously introduced ProDock ^10^ as an open-source toolkit for multi-target docking campaigns, provenance-aware execution, interaction-fingerprint postprocessing, and SQLite -backed storage. The present work extends that software layer beyond campaign organization and storage to rank-resolved multi-engine re-ranking and benchmark-driven threshold optimization. Lower-ranked poses are not necessarily uninformative: because protein-ligand binding is dynamic, a pose that the engine orders incorrectly may still be physically meaningful and may recover favorable interactions, correct binding-pocket localization, or steric compatibility.

Here, we describe the extended ProDock workflow for DiffDock and GNINA execution, rank-level descriptor calculation, and Optuna -based threshold optimization. The central design choice is to preserve multiple docking ranks and evaluate each pose independently using engine-specific scores plus additional descriptors such as binding-site occupancy, ligand-atom localization, interaction-fingerprint similarity, and steric clash counts. We benchmarked this rank-resolved re-ranking strategy on the DUDE-Z dataset, ^11^ a more stringent derivative of DUD-E^12^ designed to reduce artificial enrichment from property-matched decoys, to assess re-ranked pose quality and virtual-screening performance, followed by Optuna ^13^ threshold optimization to identify the docking score combinations that improve active-compound enrichment. The result is a reproducible framework for transforming heterogeneous docking outputs into a structured, multi-metric decision process suitable for virtual screening campaigns.

## 2 Methods

### 2.1 System Overview

ProDock is a semi-automated Python package for protein preparation, ligand preparation, docking execution, and rank-resolved post-docking analysis. The workflow combines global docking with DiffDock and local docking with GNINA, preserving multiple poses per ligand instead of reducing each docking run to a single top-ranked conformer. Each pose rank is evaluated independently with engine-specific scores and additional descriptors. For GNINA, these descriptors include empirical affinity, CNNpose, CNNaffinity, reference-based protein-ligand interaction-fingerprint similarity, steric clash counts, and an optional electrostatic solvation energy. For DiffDock, they include the confidence score, two complementary SASA localization scores, and the percentage of ligand atoms located in the binding site.

ProDock uses RDKit,^14^ PyMOL,^15^ Open Babel,^16^ and OpenMM ^17^ for structure preparation. Analysis uses MDAnalysis,^18^ ProLIF,^19^ Biopython,^20^ and APBS .^21,22^ The package writes structured intermediate files and target-specific output directories so that preparation, docking, re-ranking, and later inspection can be repeated from explicit inputs.

### 2.2 Detailed Workflow

#### 2.2.1 Automated Protein Processing and Curation

ProDock prepares receptor structures from Protein Data Bank entries or user-supplied co-ordinate files. A comma-separated input table specifies the PDB identifier, protein chains, co-crystallized ligand code, cofactors to retain, and optional residue-renumbering preferences. For each target, the PyMOL command-line interface extracts the requested chains and cofactors, removes solvent molecules and crystallization additives unless explicitly retained, and exports a receptor file for downstream processing. Co-crystallized ligands are isolated and converted to .sdf format with Open Babel . These structures can be used as reference ligands for binding-site definition and interaction-fingerprint comparisons.

Missing heavy atoms, missing residues, and protonation states are handled before docking. The preparation workflow (see Figure 1) adds hydrogen atoms at pH 7.4 and can perform an optional OpenMM -based minimization to relieve local steric strain introduced during structure repair. Receptors are parameterized with the AMBER14 force field, and backbone atoms are restrained during the initial minimization by a harmonic potential of default force constant *k* = 5.0 kcal mol^−1^ Å^−2^ anchored at the input coordinates. The default minimization stops when the energy tolerance reaches 1 kJ mol^−1^ or after 500 iterations. When solvent-phase relaxation is requested, the minimized receptor is solvated with TIP3P water in a box with 1.0 nm padding and 150 mM NaCl, with electrostatics evaluated by Particle Mesh Ewald. The final receptor is stripped of waters and neutralizing ions unless the user explicitly retains them, preserving only the prepared protein chains and selected cofactors.

**Figure 1:**
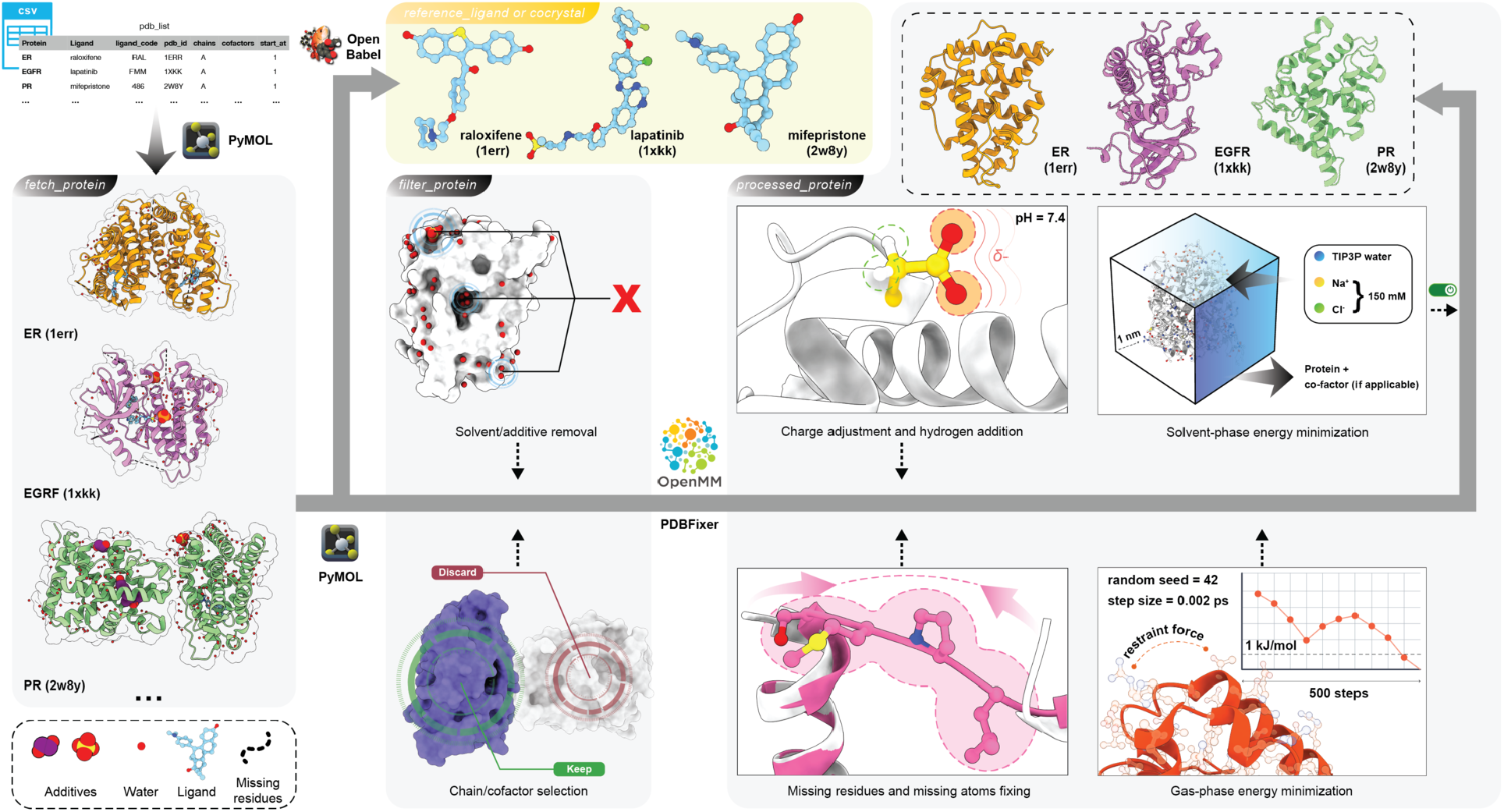
ProDock receptor preparation from structure curation through docking export.

#### 2.2.2 Automated Ligand Preparation

Ligands are prepared with RDKit from either a multi-molecule .sdf file or a .csv table containing compound names and SMILES strings. ProDock uses Chem.SDMolSupplier for .sdf input and Chem.MolFromSmiles for tabular input. Each ligand is processed independently, assigned a reproducible output filename, and written to an individual .sdf file. Invalid structures, invalid SMILES strings, embedding failures, and minimization failures are skipped and recorded in a log file.

Explicit hydrogen atoms are added with Chem.AddHs . Three-dimensional conformers are generated with AllChem.EmbedMolecule using a fixed random seed of 42, followed by geometry optimization with the standard MMFF force field (AllChem.MMFFOptimizeMolecule ), whose energy sums the usual bonded (bond, angle, torsion) and nonbonded (van der Waals, electrostatic) contributions. The ligand preparation workflow is illustrated in Figure 2.

**Figure 2:**
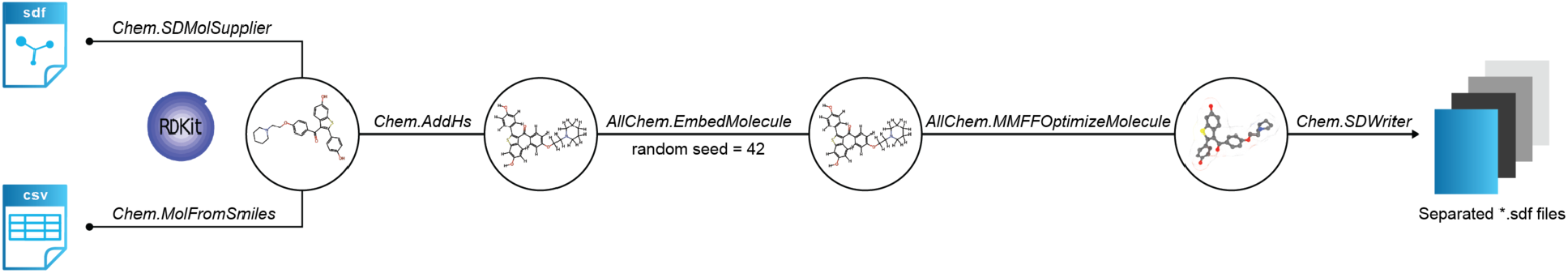
ProDock ligand preparation from .sdf or .csv input to minimized 3D structures.

#### 2.2.3 Automated Docking Process

ProDock provides batch modules for all-by-all receptor-ligand docking with DiffDock and GNINA . The DiffDock module takes directories of receptor .pdb files and ligand .sdf files, generates target-specific output folders, and allows user control over the model directory and the number of generated samples per complex. The GNINA module performs local docking with either automatic box generation from a reference ligand through GNINA’s autobox option or manually specified box centers and dimensions. Default local-docking settings use a random seed of 42, exhaustiveness of 32, resulting in 10 generated poses per ligand. Multi-conformer GNINA outputs are split into rank-specific pose files before analysis.

#### 2.2.4 Automated Docking Analysis

##### Rank organization and binding-site definition

ProDock treats each generated pose rank as an independent analysis object. If *n* poses are produced, the workflow creates rank-specific files for *i* = 1, …, *n*, with one pose per ligand in each rank directory. The downstream descriptors are engine specific: DiffDock filtering emphasizes localization in the binding site, whereas GNINA filtering emphasizes local binding modes, interaction recovery, energetic scores, and steric feasibility.

Binding-site residues are defined either from a reference ligand, typically the crystallographic ligand, or from a user-supplied residue list. For reference-based site definition, residues within a user-defined cutoff distance *d*_cut_ of the reference ligand are selected with PyMOL . The default cutoff is 5 Å. Let the protein be represented by residues

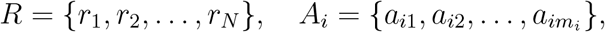

where *A*_*i*_ is the atom set for residue *r*_*i*_. Let the reference ligand atoms be

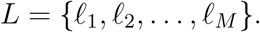

The Euclidean distance between protein atom *a*_*ij*_ and ligand atom *ℓ*_*k*_ is

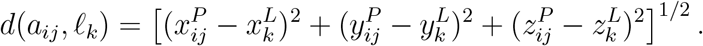

A protein residue *r*_*i*_ is assigned to the binding site if

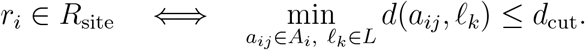

##### DiffDock localization descriptors

For DiffDock poses, ProDock measures how well the ligand localizes to the binding region. With PyMOL dot_solvent enabled, the workflow estimates solvent accessible surface areas for the isolated ligand (*A*_lig_), isolated binding site (*A*_site_), and ligand plus site complex (*A*_complex_). Following the software implementation, the residual SASA score stored as Unoccupied is

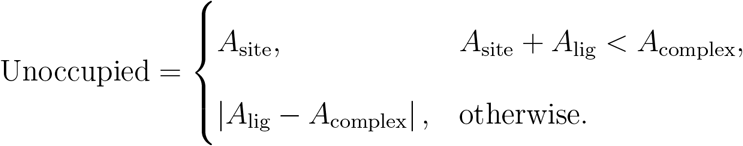

The corresponding normalized score is

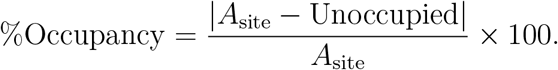

For the usual numerical regime *A*_lig_ ≤ *A*_complex_ ≤ *A*_lig_ + *A*_site_, the residual score reduces to Unoccupied = *A*_complex_ − *A*_lig_ = *A*_site_ − (*A*_lig_ + *A*_site_ − *A*_complex_). The term in parentheses is the total buried SASA across both molecular surfaces. These quantities are therefore operational SASA localization scores rather than a pocket volume or a direct measure of the protein side contact area. Lower Unoccupied and higher %Occupancy indicate stronger spatial overlap. Because the two scores are deterministically related for a fixed target, the default filter retains only %Occupancy.

The additional DiffDock localization descriptor is the percentage of ligand atoms located inside the binding region. For a ligand pose *D* = *{q*_1_, *q*_2_, …, *q*_*Q*_*}*, atom *q* is considered inside the site if its distance to any atom in *R*_site_ is less than or equal to *d*_cut_:

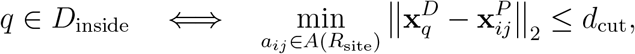

where *A*(*R*_site_) denotes all atoms belonging to the selected binding-site residues. The atom-localization percentage is

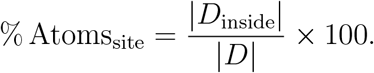

##### GNINA interaction and energy descriptors

For GNINA poses, ProDock compares the docked pose with the reference ligand using ProLIF interaction fingerprints. Two similarity scores are reported. Type 1 similarity is strict: a docked pose must recover the same residue and interaction type as the reference. Type 2 similarity is residue-based: a docked pose is credited for contacting the same residue even if the interaction type differs. Let *F* ^(1)^ be the set of residue-interaction features,

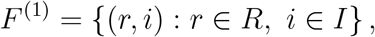

where *R* is the residue set and *I* is the set of interaction types detected by ProLIF . With 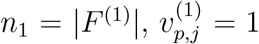 if pose *p* contains feature *j* and 0 otherwise. For pose *p* and reference ligand ref, type 1 similarity is

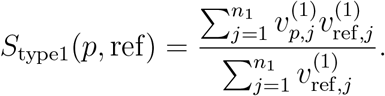

For type 2 similarity, interaction types are collapsed to residue-level contacts. Let

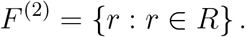

With *n*_2_ = |*F* ^(2)^|, the residue-level fingerprint is obtained from the type 1 vector by

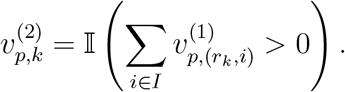

The type 2 similarity score is

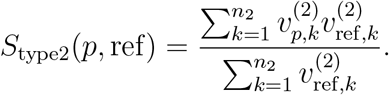

The denominators count the reference features. If a reference contains no detectable interaction features for a given type, the corresponding similarity is reported as missing rather than forced to be zero.

ProDock can optionally estimate ligand electrostatic solvation energy with APBS after structure conversion with Open Babel . This descriptor was not computed for the DUDE-Z benchmark and was excluded from the analyses reported here.

##### Rank-level filtering and pose selection

Rank-level metrics are merged into target-specific summary .csv files. A rank is considered satisfied when all required criteria pass. For DiffDock, the default final label for rank *i* is

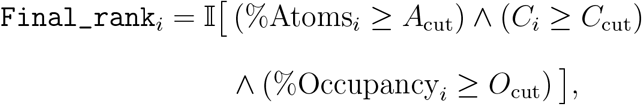

where *C*_*i*_ is the DiffDock confidence score, %Occupancy_*i*_ is the binding-site occupancy percentage, and %Atoms_*i*_ is the percentage of ligand atoms inside the site. The default thresholds are *A*_cut_ = 80.0, *O*_cut_ = 20.0, and *C*_cut_ = −1.5.

For GNINA, the default final label is

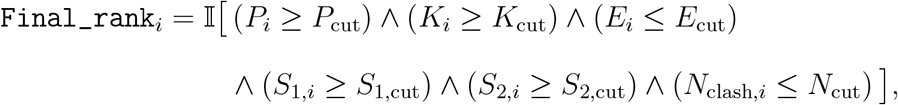

where *P*_*i*_ is CNNpose, *K*_*i*_ is CNNaffinity, *E*_*i*_ is empirical docking affinity, *S*_1,*i*_ and *S*_2,*i*_ are type 1 and type 2 similarities, and *N*_clash,*i*_ is the steric clash count. The default thresholds are *P*_cut_ = 0.3, *K*_cut_ = 0, *E*_cut_ = 0, *S*_1,cut_ = 0, *S*_2,cut_ = 0, and *N*_cut_ = 2. The steric clash count is calculated using a default clash-distance threshold of 1.5 Å, where any protein-ligand atom pair below this distance is counted as a clash.

With *F*_*i*_ denoting the binary satisfaction label for rank *i*, the number of satisfied ranks ^is 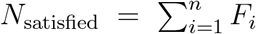^, and a compound is retained when *N*_satisfied_ *>* 0. The reported best_satisfied_rank is the first passing rank, *r*_best_ = min*{i* ∈ *{*1, 2, …, *n}* : *F*_*i*_ = 1*}*. Thus, when several ranks pass, the earliest acceptable pose is selected. If no rank passes, the compound is rejected by that filter set. Algorithm 1 summarizes this procedure. The generalized acceptance predicate and its conversion into a compound-level ranking are provided in Supporting Section S1.1.

###### Algorithm 1 Rank-resolved filtering and pose selection in ProDock

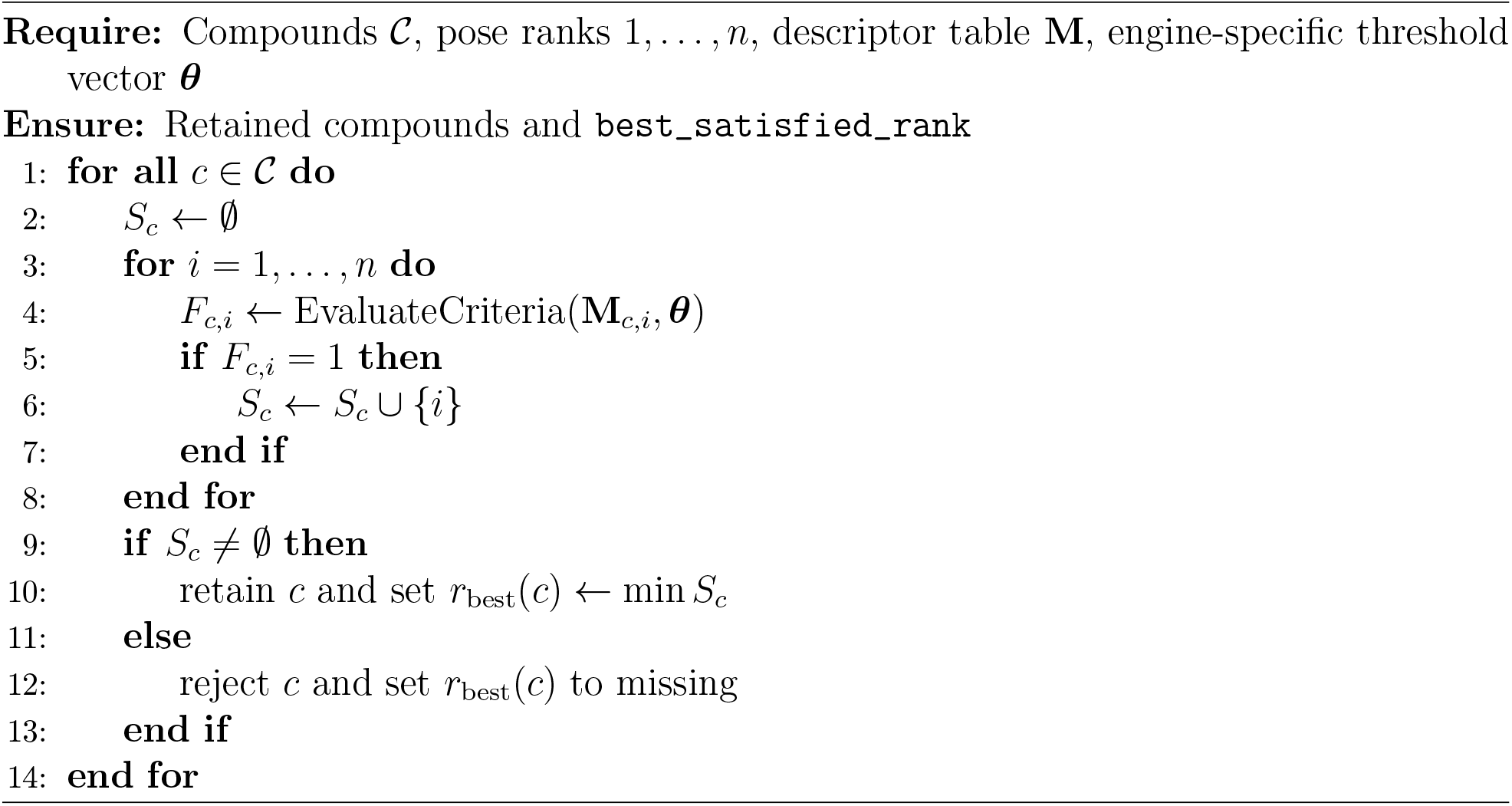

The overall pipeline of the docking process, analysis, pose selection, and re-ranking in ProDock is summarized in Figure 3.

**Figure 3:**
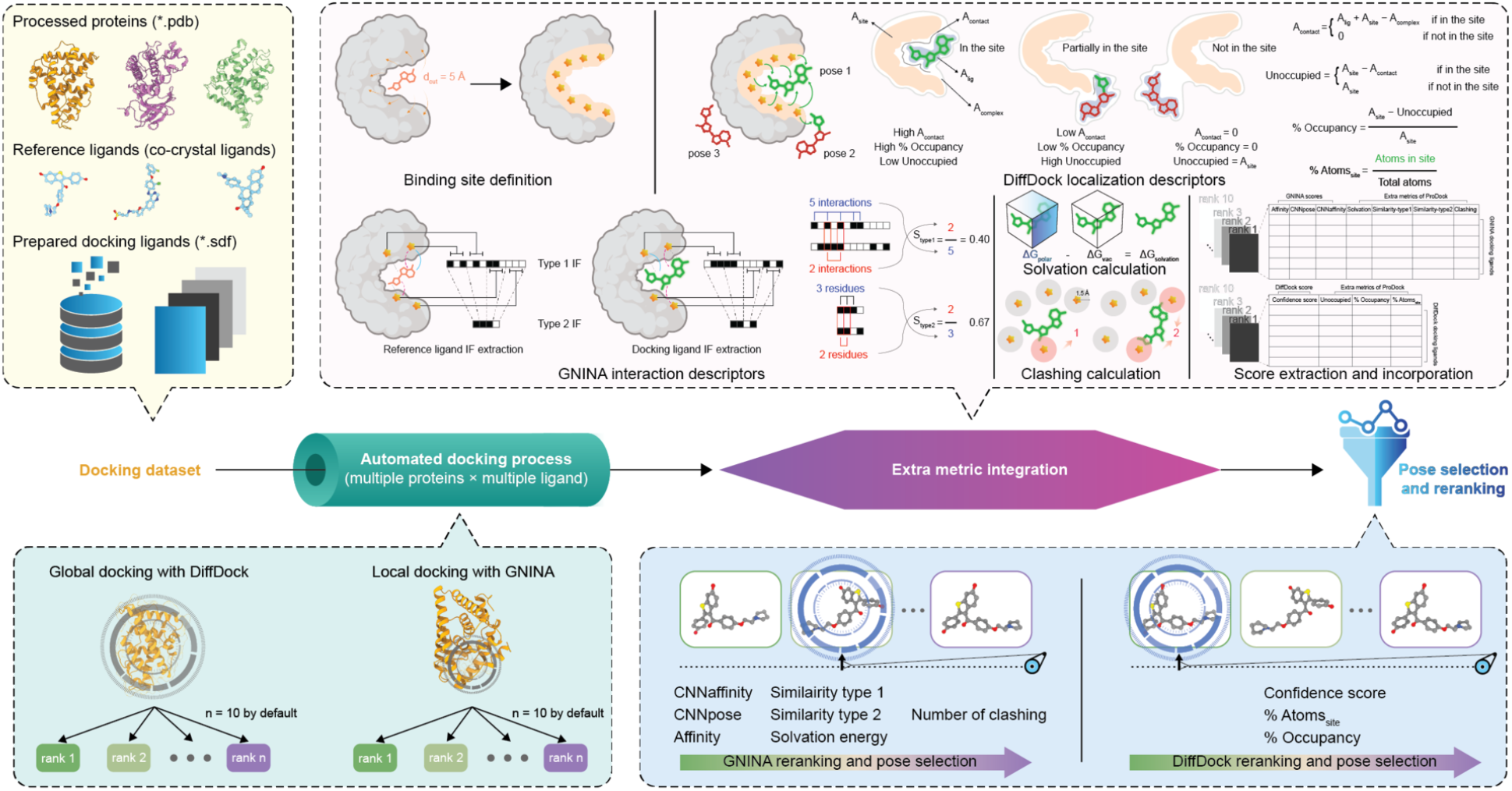
ProDock workflow for preparation, multi-engine docking, pose-level analysis, filtering, and re-ranking.

ProDock also provides optional property reporting through ADMETLab 3.0 .^23^ This module was not used in the DUDE-Z benchmark, and its workflow is described in Supporting Section S2.

### 2.3 Rank-Resolved Re-Ranking

The per-pose filter of Section 2.2.4 decides whether a pose is acceptable, but virtual screening requires a compound ranking. For a compound with pose ensemble *p*_1_, …, *p*_*n*_, each pose is accepted or rejected by the conjunction of threshold criteria *F* defined above. A retained compound is represented by its *first* accepted pose, which preserves the engine’s ordering among acceptable poses. That pose is scored by empirical affinity, CNNaffinity, or the composite CNNxAffinity, which is the product of CNNpose and CNNaffinity . The deposited branch assigns no score to compounds with no accepted rank and omits them from the screened score table. The separate optimization script used to generate the archived benchmark metrics was not deposited, so any additional fallback behavior in that analysis cannot be independently verified. The conservative formalization supported by the available implementation is given in Supporting Section S1.1 (Algorithm S1).

### 2.4 DUDE-Z Docking and Optimization

#### 2.4.1 DUDE-Z Docking

DUDE-Z ^11^ was used to evaluate the dual global-local docking and rank-resolved re-ranking workflow. The benchmark comprises 43 pharmaceutically relevant protein targets with experimentally supported active ligands and property-matched decoys. Compared with simple random decoys, DUDE-Z decoys are designed to resemble active compounds in physico-chemical descriptors and to match net charge at physiological pH, making enrichment less dependent on trivial ligand-property differences.

For each target, actives and decoys were docked globally with DiffDock and locally with GNINA . The GNINA settings were --seed42, --cnn_scoringrescore, --exhaustiveness32, and --num_modes10 . The co-crystal ligand defined --autobox_ligand, with --autobox_a dd5 . It also defined --flexdist_ligand, with --flexdist5 .

DiffDock used its default_inference_args.yaml settings. The resulting ranks were processed with the same analysis modules used in prospective screening. Inputs comprised:

- 3D protein structural information deposited in the rec.crg.pdb files.
- 3D structural information on the co-crystallized ligand deposited in the xtal-lig.pdb files.
- 2D structural information on known ligands and decoys, deposited in ligand.smi and decoy.smi, respectively, in the form of isomeric SMILES notations, with protonation states calculated for the physiologically relevant pH.

These data were converted into receptor .pdb files and ligand .sdf files within the directory structure expected by ProDock .

#### 2.4.2 Optuna Optimization

Docking scores and ProDock descriptors from GNINA and DiffDock were aggregated by unique compound identifiers to ensure paired data availability. For each target independently, its compounds were partitioned into a training set (80%) and a held-out test set (20%), stratified by binary activity label (Table S3 of the Supporting Information gives the per-target split sizes). Threshold optimization was likewise performed separately for every target. Each of the three scoring configurations, empirical affinity, CNNaffinity, and CNNxAffinity (Section 2.3), was optimized against each of the three objectives, giving nine scoring*×*metric studies per target.

Each Optuna study ran 400 trials with the default Tree-structured Parzen Estimator sampler and no pruner, maximizing the training-split objective. Up to eight thresholds were treated as continuous variables sampled uniformly between the empirical minimum and maximum of the corresponding descriptor over the ten-pose pool of each molecule. When the composite CNNxAffinity score was used for ranking, CNNpose was removed from the threshold pool to avoid using the same quantity as both a ranking score and a filter. According to the archived study configuration, the optimized threshold vector ***θ***^∗^ was frozen and applied once to the held-out test split. The generating optimization script was not deposited, and the available screening branch omits compounds that pass no rank. Exact equality of the baseline and optimized compound sets therefore cannot be independently verified. The reported metric differences are treated as comparisons of archived pipeline outputs rather than as confirmed paired measurements over identical compounds. The full study configuration and the resulting per-target threshold vectors are given in Section S3 and Table S13 of the Supporting Information. The conservative optimization formalism is stated in Section S1.2 of the Supporting Information.

### 2.5 Evaluation Metrics

Screening performance is reported using the archived ROC-AUC, PR-AUC, and LogAUC outputs. LogAUC is a logarithmically weighted ROC summary that emphasizes the low false positive rate region. Its generating code and lower integration cutoff were not deposited, so the exact numerical convention cannot be verified and comparisons are restricted to values carrying the same archived metric label (Supporting Section S1.3). Because the benchmark is strongly class imbalanced (2,376 actives among 136,759 docked compounds, with a median decoy-to-active ratio of 56:1 per target), PR-AUC is reported alongside ROC-AUC. ROC-AUC is insensitive to the large decoy excess, whereas PR-AUC responds directly to the precision of the top of the ranked list and is the more faithful indicator of practical enrichment.

Rank-level active-decoy discrimination for a *single* descriptor, reported per target and per pose rank in Section 3.1 and Section 3.2, was quantified with a separation coefficient, defined as the complement of the overlapping coefficient of the active and decoy value distributions. With the two classes histogrammed over their common range into *B* = 30 unit-area bins of width *w* and densities 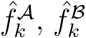, the coefficient is

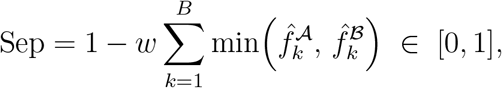

equal to 0 when the two distributions coincide and 1 when they are disjoint. Being a density-overlap rather than a rank statistic, it is sensitive to outliers and to the binning. Descriptor columns are screened for sentinel values before it is computed, as detailed in Section S1.3 of the Supporting Information.

## 3 Results and Discussion

### 3.1 GNINA Docking Scores across All Ranks and DUDE-Z Benchmark Targets

All GNINA metrics discriminated actives from decoys best at high pose ranks, but degraded at different rates: CNNaffinity was most rank-robust (mean separation 0.578 at rank 1 versus 0.518 at rank 10), whereas Similarity type 1 declined most (0.376 versus 0.279). Target effects were larger than rank effects (Figure 4D): mean separation ranged from about 0.8 for PUR2, XIAP, TRYB1, and FA10 to about 0.2 for ACES and ANDR, compared with a rank 1 to rank 10 decay of 0.05 to 0.10. This variation motivates multi-metric rather than single-score pose selection.

**Figure 4:**
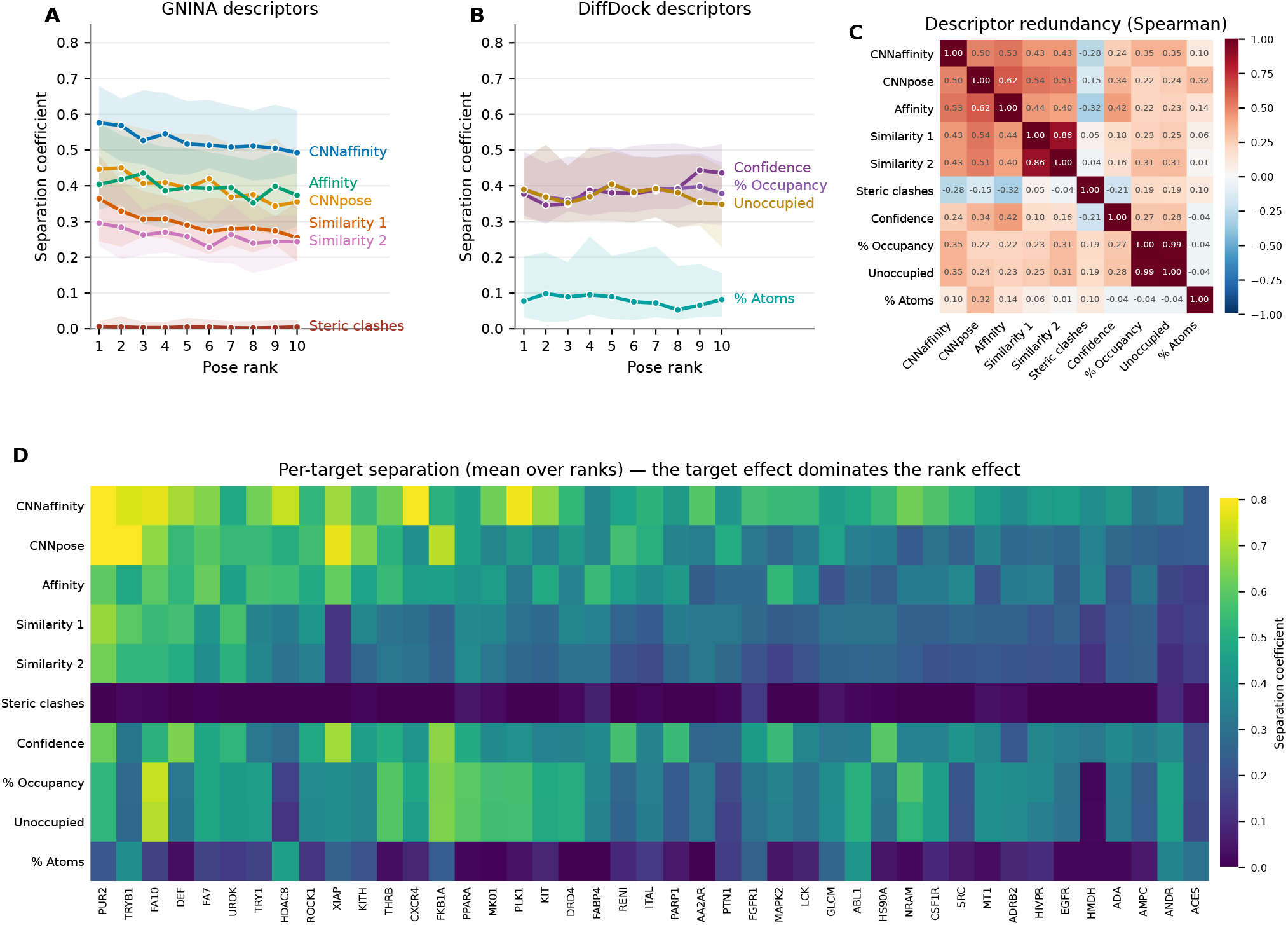
Separation of actives and decoys across 43 DUDE-Z targets. **(A**,**B)** Median separation by pose rank for GNINA and DiffDock . shading is the interquartile range. **(C)** Descriptor correlations across combinations of targets and ranks. **(D)** Rank-averaged separation by target and descriptor. Solvation energy was not computed. Numerical values are in Supporting Tables S5 through S9.

Many targets had extreme positive Affinity values, consistent with unfavorable conformations (Figure S1). Rank-averaged separation exceeded 0.55 for FA7, XIAP, PUR2, FA10, TRY1, and HDAC8 but was below 0.22 for ACES, ANDR, ADA, GLCM, and MT1.

CNNpose assigned higher scores to actives than to decoys, particularly at early ranks, and showed the steepest rank dependence of the three GNINA scores (Figure 4). Its mean separation coefficient fell from 0.4712 at rank 1 to 0.3967 at rank 10, a decrease that was significant across the 43 targets (paired Wilcoxon signed-rank test, *p* = 2.0 *×* 10^−5^). This behavior is consistent with CNNpose being trained to recognize pose quality rather than binding affinity, and it makes CNNpose a natural rank-quality criterion to combine with affinity-like scores during optimization.

By contrast, CNNaffinity gave the highest separation coefficient of any metric at every rank (Figure 4 and Figure S3) and was the most rank-robust, retaining a mean separation of 0.5181 even at rank 10. CNNaffinity therefore discriminates actives from decoys largely independently of whether the pose itself is well placed, which makes it a strong ranking score but a weak pose filter. This complementarity motivates combining it with pose-quality descriptors rather than using it alone.

The similarity metrics behaved in parallel (Spearman *ρ* = 0.86) but type 1, which requires the interaction type as well as residue to match, separated the classes more strongly. Its mean separation declined by 25%, from 0.3758 to 0.2791 (paired Wilcoxon test, *p* = 4.5 *×* 10^−6^). The active and decoy gap narrowed from 0.1107 to 0.0765 (Figures S4 and S5). Type 2 tolerates more binding modes but loses some discrimination. Both are reference anchored and may enrich ligands resembling the co-crystal ligand, a limitation addressed below.

### 3.2 DiffDock Localization Descriptors across All Ranks and DUDE-Z Benchmark Targets

Localization behavior differed between the two engines. The DiffDock separation coefficients are shown in Figure 4, with distributions across ranks and targets in Supporting Figures S6 through S8.

Two observations follow. First, unlike every GNINA score, the DiffDock descriptors were essentially rank independent. The mean %Atoms separation was statistically indistinguishable between rank 1 and rank 10 (0.13 versus 0.13, paired Wilcoxon signed-rank test, *p* = 0.89), and %Occupancy changed only marginally (0.389 versus 0.374, *p* = 0.08). DiffDock confidence ordering therefore carries little information about how well a pose localizes in the reference binding site, which is consistent with DiffDock performing a global, blind search rather than a site-restricted one.

Second, %Atoms discriminated actives from decoys only weakly (mean separation 0.13, median 0.08, versus 0.55 for CNNaffinity ), and its separation coefficient was uncorrelated with those of the GNINA scores (Spearman *ρ* ≤ 0.14 against Affinity and CNNaffinity ). This is expected, because DUDE-Z decoys are property matched to the actives and are of comparable size: a decoy is about as likely as an active to place most of its atoms inside a pocket. The descriptor is thus best used as a pose-validity gate that removes grossly mislocalized DiffDock poses rather than as an enrichment score, which is how it is applied in the default filter (*A*_cut_ = 80).

We further note that %Occupancy (Figure S7) and Unoccupied (Figure S8) are inversely related by construction at the pose level. Their unsigned active versus decoy separation coefficients consequently carried near identical information (Spearman *ρ* = 0.99 across all 430 target and rank combinations). They should be treated as one descriptor, and only one of the two need be retained in a filter.

### 3.3 Optuna Optimization

The archived held-out tables report higher optimized values than rank 1 across most scoring configurations. These differences are descriptive because identical baseline and optimized compound membership cannot be verified from the deposited code.

Both CNN-derived scoring functions had higher archived values across ROC-AUC, PR-AUC, and LogAUC (Figure 5A and B). The largest difference was for CNNaffinity PR-AUC, from 0.197 to 0.294 (+0.097, 49% relative). Its LogAUC changed from 0.708 to 0.763, and CNNxAffinity ROC-AUC changed from 0.731 to 0.771. Across scoring functions, PR-AUC differences were +0.068 for Affinity, +0.097 for CNNaffinity, and +0.041 for CNNxAffinity . ROC-AUC differences were smaller and score dependent, ranging from −0.012 for Affinity (0.770 to 0.758) to +0.040 for CNNxAffinity . Within the archived outputs, the largest positive differences occurred for early enrichment and CNN-based scores. Per-target analysis across the 43 DUDE-Z targets showed substantial variation between proteins, as shown in Figure 5C. The number of positive archived differences ranged from 22 out of 43 for Affinity ROC-AUC to 29 out of 43 for Affinity LogAUC. The lower end of this range is indistinguishable from chance and should not be over-interpreted. The upper end (29/43, 67%) shows that a majority of the archived target comparisons were positive for the LogAUC objective. The distribution of per-target differences is broad rather than uniformly positive.

**Figure 5:**
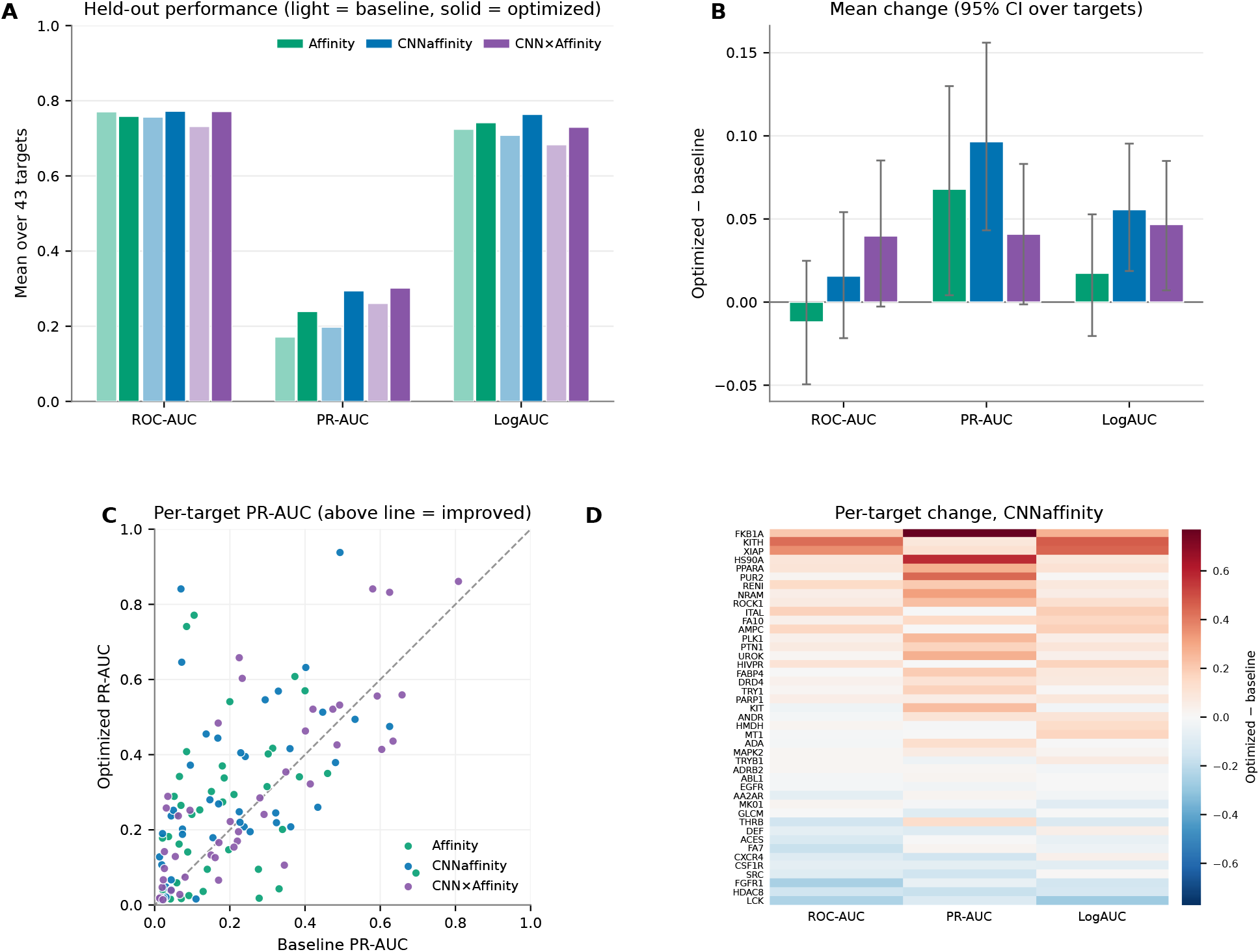
Archived held-out metric values for optimized re-ranking versus rank 1 across 43 targets. **(A)** Mean baseline and optimized metrics for three scoring functions. **(B)** Mean differences with 95% bootstrap confidence intervals. **(C)** Per-target PR-AUC. **(D)** Pertarget metric changes for CNNaffinity .

Baseline-to-optimized PR-AUC correlation was strongest for CNNxAffinity (Pearson *r* = 0.81), weakest for Affinity (*r* = 0.24), and intermediate for CNNaffinity (*r* from 0.56 to 0.73 across metrics), indicating more consistent archived target-level patterns for CNN-derived scores.

The per-protein deltas (Figure 5D) show substantial positive differences for a subset of targets, notably FKB1A, PUR2, HS90A, and KITH, alongside smaller or negative differences elsewhere.

Because thresholds were optimized independently for every target, the selected values form a distribution across targets rather than a single universal setting, and their spread indicates how consistently the optimizer relied on each criterion (Figure 6). The interaction-similarity and CNNpose thresholds were broadly distributed, consistent with target-specific tuning, whereas the DiffDock %Atoms threshold was driven to its maximum of 100% in 34 of 43 targets. This saturating behavior shows that %Atoms functioned as a near-binary pose-validity gate rather than a graded filter, in keeping with its weak standalone active-decoy separation (Section 3.2). The full per-target threshold vectors are provided in the Supporting Information (Table S13).

**Figure 6:**
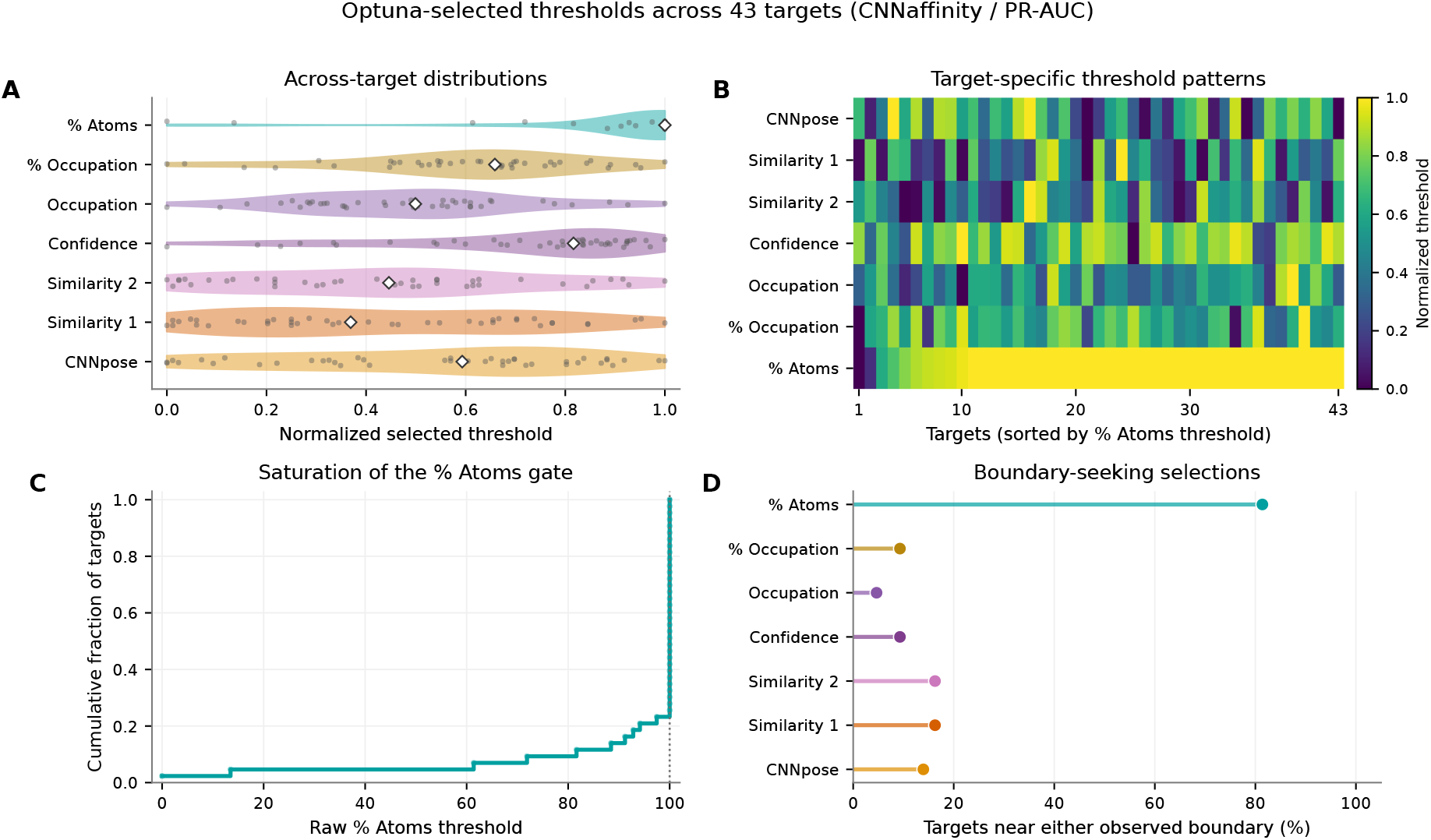
Optuna -selected thresholds across 43 targets for CNNaffinity /PR-AUC. **(A)** Normalized distributions and medians. **(B)** Target-level normalized thresholds. **(C)** Raw %Atoms empirical cumulative distribution. **(D)** Targets within 5% of an observed descriptor boundary.

In contrast, LCK and THRB showed consistent decreases across several scoring configurations. LCK showed reductions in ROC-AUC and LogAUC, while THRB also decreased substantially for both metrics. HDAC8 and FGFR1 showed similar but smaller negative trends. Two explanations are compatible with these observations, and we cannot presently distinguish between them. The first is statistical: the benchmark provides a median of only 49 actives per target (range 23-95), so a 20% held-out split leaves fewer than ten test actives for several targets, and per-target PR-AUC estimated from so few positives is intrinsically noisy. The second is mechanistic: re-ranking may be less reliable for targets with highly flexible binding sites, promiscuous ligand recognition, or target-specific decoy-design biases not captured by the optimization framework. Distinguishing these requires resampling-based confidence intervals on the per-target deltas, which we identify as necessary future work rather than asserting a mechanism the present data cannot support. One notable example was XIAP, where Affinity PR-AUC decreased substantially while CNNxAffinity ROC-AUC increased strongly. The archived comparisons are consistent with target and score dependent effects, but confirmation requires recomputation over a fixed compound set.

### 3.4 Structural Basis of Pose Recovery

To investigate the structural mechanism associated with the reported enrichment differences, we examined the docked poses for a subset of rescued targets for which an experimental co-crystal structure was available: neuraminidase (NRAM, PDB 1b9v), the X-linked inhibitor of apoptosis protein (XIAP, PDB 3hl5), and HIV-1 protease (HIVPR, PDB 1xl2). For each target we compared, across the held-out test-set active compounds, the default rank-1 pose against the pose selected by Optuna -optimized re-ranking, using the co-crystallized ligand as the native reference. All poses and reference ligands were analyzed in the original docking coordinate frame without superposition.

Across the 18 rescued test-set actives in these three targets, re-ranking moved the selected pose closer to the native binding pose than the rank-1 pose (Figure 7). Per-compound values are reported in Table S14 of the Supporting Information. On average, the fraction of ligand heavy atoms occupying the native binding volume (defined as the region within 2 Å of any co-crystal ligand atom, which differs from the %Occupancy descriptor above so as to preserve the true extent of the binding region) increased by 0.12, the distance between the pose and native-ligand centroids decreased by 0.73 Å, and the residue-level interaction-fingerprint similarity to the co-crystal ligand increased by 0.17. Improvement was observed in 14 of 18 compounds for binding-site occupancy, 15 of 18 for centroid proximity, and all 18 for interaction similarity (Figure 7A to C), indicating that the effect was not restricted to isolated cases. The re-ranking selected poses at ranks 2 through 10 rather than rank 1 (Figure 7D), confirming that the improvement came from genuinely reordering the pose ensemble.

**Figure 7:**
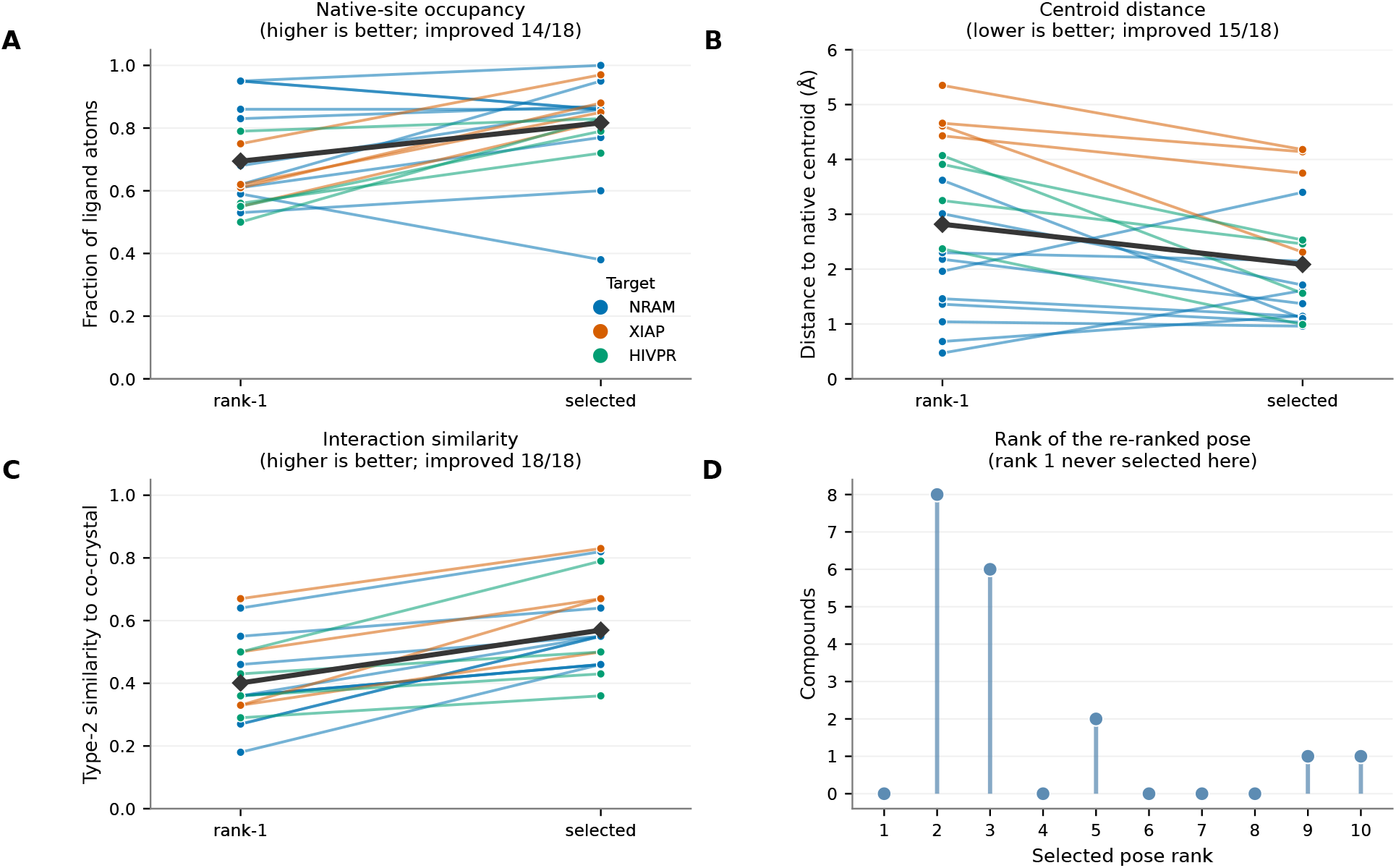
Structural rescue for 18 held-out actives (NRAM in blue, XIAP in orange, and HIVPR in green). Lines connect rank 1 to the selected pose. Black diamonds show means. **(A)** Native-site occupancy. **(B)** Native-centroid distance. **(C)** Type-2 co-crystal interaction similarity. **(D)** Selected rank.

The overlays in Figure 8 show two distinct structural mechanisms. For XIAP and HIVPR, re-ranking produced a spatial rescue. The rank-1 poses were substantially displaced from the native site (CHEMBL375784 in XIAP, centroid distance 4.6 Å, and CHEMBL119970 in HIVPR, 2.4 Å). The re-ranked pose relocated into the native binding volume (2.3 Å and 1.0 Å, respectively), increasing occupancy of the native site from 0.55 to 0.82 for XIAP and from 0.56 to 0.72 for HIVPR. For NRAM, re-ranking produced an interaction rescue. The rank-1 poses were already localized within the binding site (CHEMBL254679, centroid distance 1.4 Å), so the improvement arose not from gross repositioning but from selecting a pose that recovered native protein-ligand interactions, increasing the interaction-fingerprint similarity from 0.27 to 0.55.

**Figure 8:**
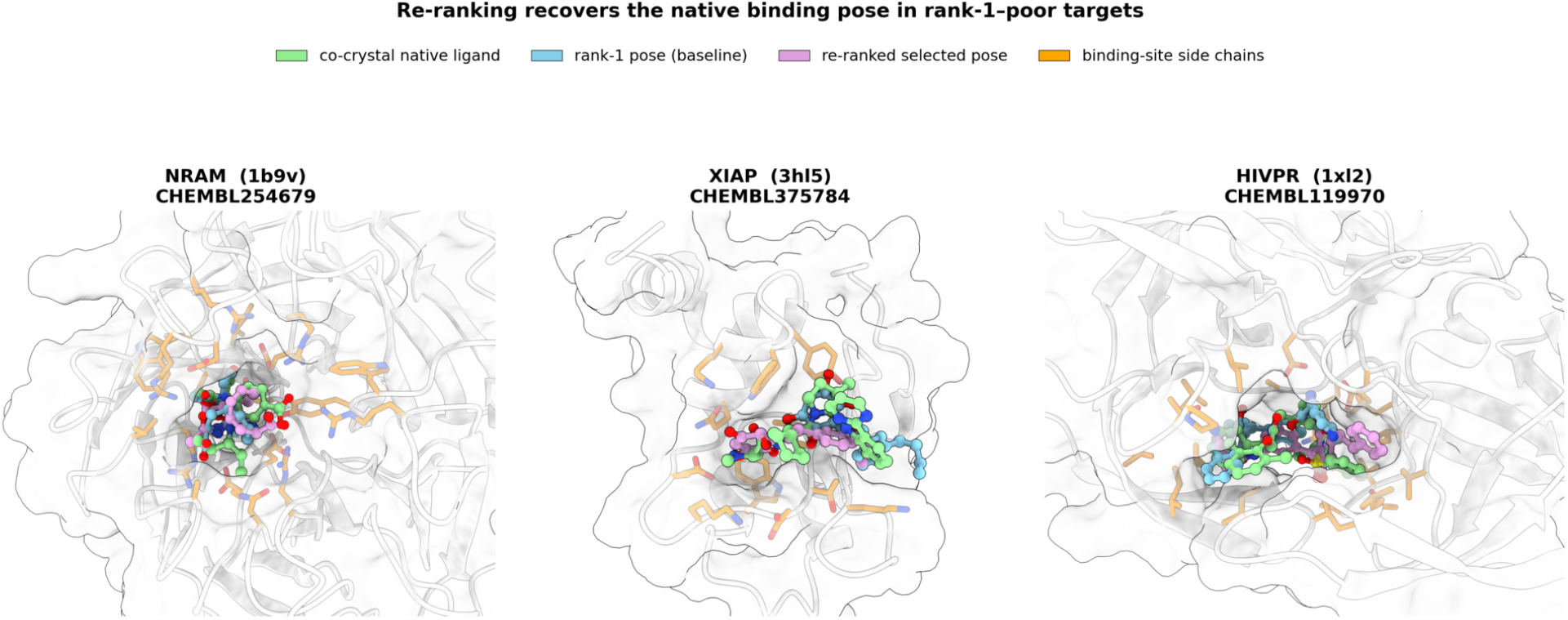
Structural overlays of rank 1, the selected pose, and the co-crystal ligand.

The interaction-level analysis confirmed this recovery (Figure 9). Using ProLIF interaction fingerprints computed against the co-crystal ligand, the number of native contacts recovered by the selected pose exceeded that of the rank-1 pose in all three representative compounds: from 2 to 7 of 9 native contacts for NRAM, 3 to 4 of 7 for XIAP, and 7 to 11 of 13 for HIVPR. In several cases re-ranking restored specific native interactions absent from the rank-1 pose. For NRAM CHEMBL254679, the rank-1 pose failed to form the single native hydrogen bond to ARG216 and instead engaged peripheral residues, whereas the re-ranked pose recovered this hydrogen bond along with the surrounding native hydrophobic contacts. For XIAP CHEMBL375784, the re-ranked pose recovered a native contact to GLN67 that was absent in the rank-1 pose.

**Figure 9:**
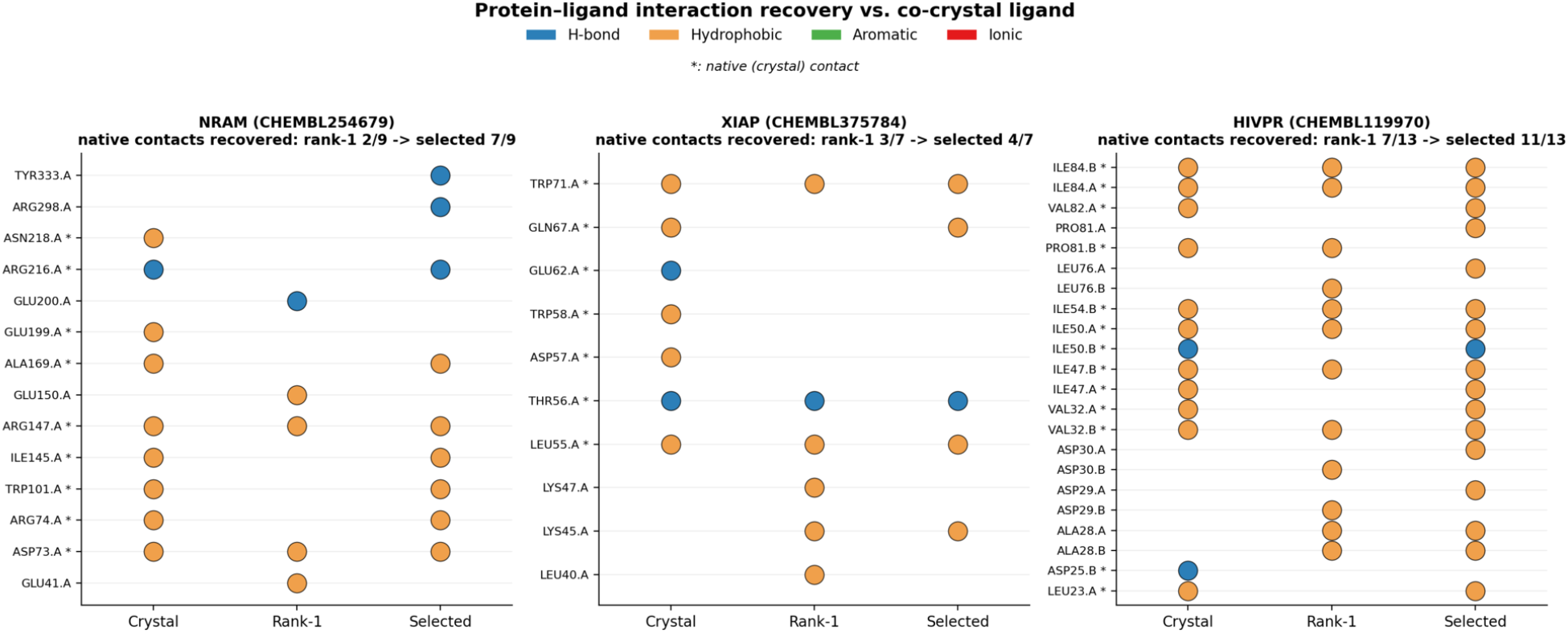
Native contacts between proteins and ligands recovered by rank 1 and the selected poses.

Because the interaction-fingerprint similarity is one of the threshold criteria used during re-ranking, these structural observations are best interpreted as a mechanistic illustration of pose selection rather than as an independent validation. The quantitative screening evidence remains the archived enrichment metrics reported above, subject to the evaluation set limitation. Nevertheless, the concurrent improvement in binding-site occupancy and centroid proximity, which are geometric measures not used as optimization criteria, indicates the re-ranked poses were genuinely more native-like. This supports the interpretation that the reported enrichment differences can coincide with recovery of physically meaningful binding modes.

## 4 Discussion and Conclusion

This work extends ProDock from campaign organization and database storage to benchmark-optimized rank-resolved re-ranking. It integrates complementary information otherwise fragmented across preparation scripts, engine-specific score files, and manual post-processing. Automated preparation, global DiffDock and local GNINA docking, and rank-resolved metrics form a reproducible sequence of inputs, intermediate files, and decision tables.

On the DUDE-Z benchmark, the archived outputs showed higher values most clearly for the CNN-derived scoring configurations and for PR-AUC and LogAUC. The differences were not uniform. Some targets had lower values after optimization, and the same target could respond differently to different scoring functions. These results motivate target-aware calibration and transparent reporting of both positive and negative target-level outcomes rather than a single universal threshold set.

Several caveats bound these conclusions. The optimization and AUC generating script was not deposited. The available screening branch omits compounds that pass no rank, so identical baseline and optimized compound membership and the exact LogAUC convention cannot be independently verified. The ranking scores are all GNINA -derived and the DiffDock descriptors enter only the multi-metric filter as optimized thresholds (Table S13), so pose selection is cross-engine, but the contribution of each engine is not isolated. The saturating %Atoms threshold suggests the localization criteria act mainly as validity gates. The interaction-similarity criteria are anchored to each target’s co-crystal ligand, which actives resemble more than decoys do, so their contribution is partly circular and may not transfer to novel chemotypes or targets lacking a co-crystal structure. The held-out split is over compounds within each target and stratified only by activity, so analog-rich series appear on both sides and the benchmark concerns unseen ligands within these 43 targets rather than unseen proteins. A scaffold-based split would be stricter. Each per-target value is a single-split point estimate, and eight thresholds fitted against a median of roughly forty training actives leave the per-target optimization exposed to overfitting that the held-out evaluation limits but does not remove. Finally, pose quality is judged by occupancy, centroid distance, and interaction recovery rather than symmetry-corrected RMSD or physical-validity checks, and the structural analysis rests on 18 compounds across three targets. Depositing the generating script, reporting retained counts, recomputing metrics over a fixed compound set, using a scaffold split, repeating the split, and adding RMSD and PoseBusters validation are the natural next steps.

The current implementation is therefore best viewed as a practical, extensible framework rather than a universal replacement for docking-engine scores. More broadly, the results support rank-resolved, multi-metric postprocessing as a transparent layer between docking-engine output and final compound selection.

## Acknowledgement

This work has received support from the Korea International Cooperation Agency (KOICA) under the project entitled “Education and Research Capacity Building Project at University of Medicine and Pharmacy at Ho Chi Minh City,” conducted from 2025 to 2026 (Project No. 2021-00020-3).

## Supporting Information Available

Extended re-ranking formalism (Section S1). Optional ADMET reporting (Section S2). Software and Optuna study configuration (Section S3). Benchmark composition and train/test split (Section S4 and Table S3). Score distributions for all ProDock descriptors across all pose ranks and targets (Figures S1 through S8). Per-target and per-rank separation coefficients. Per-target held-out ROC-AUC, PR-AUC, and LogAUC before and after optimization (Tables S10 through S12). Per-target optimized thresholds (Table S13). Structural rescue comparison (Table S14).

## Data and Software Availability

Benchmark data and derived result tables are available at https://github.com/Medicine-Artificial-Intelligence/ProDock/tree/main/Project/benchmark. The source code used in this study is available at https://github.com/Medicine-Artificial-Intelligence/ProDock/tree/main.

## Notes

The authors declare no competing financial interest.

## TOC Graphic

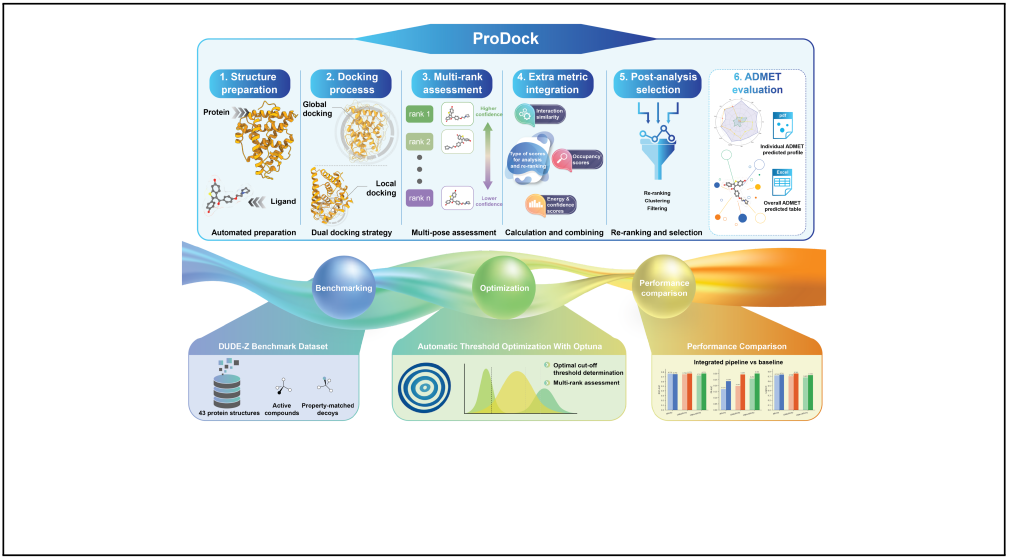

