## Supplementary material for "Rank-Resolved Multi-Engine Docking and Optuna-Optimized Re-Ranking with ProDock for Virtual Screening": SI

### S1 Re-Ranking Formulation

This section gives the formal statement of the rank-resolved re-ranking rule, the compound-level score it induces, and the threshold-optimization procedure, expanding the compact description in the Methods of the main text.

#### S1.1 Re-ranking framework

The filtering rules of the main text select a pose; virtual screening, however, requires a total order on compounds. We therefore state explicitly how a per-pose acceptance predicate is converted into a compound-level ranking, since the two objects are not interchangeable and the conversion affects every reported metric.

**Poses, descriptors, and criteria.** Let $\mathcal{C}$ be the compound library and let each compound $c \in \mathcal{C}$ be docked into a fixed receptor, yielding an ordered pose ensemble

$$\mathcal{P}(c) = (p_{c,1}, p_{c,2}, \ldots, p_{c,n}),$$

where the index $i$ is the rank assigned by the docking engine and $n = 10$ throughout this work. A descriptor map

$$\Phi : \; p_{c,i} \;\longmapsto\; \mathbf{m}_{c,i} = \left(m_{c,i}^{(1)}, \ldots, m_{c,i}^{(d)}\right) \in \mathbb{R}^d$$

assigns to every pose the engine-specific score vector of the Methods. Each descriptor $t \in \{1, \ldots, d\}$ is paired with a threshold $\theta^{(t)}$ and a direction $\sigma^{(t)} \in \{+1, -1\}$, where $\sigma^{(t)} = +1$ encodes a lower bound (larger is better, e.g. `CNNaffinity`) and $\sigma^{(t)} = -1$ an upper bound (smaller is better, e.g. the steric clash count). The acceptance predicate for a single pose is then the conjunction

$$F(\mathbf{m}_{c,i};\boldsymbol{\theta}) = \prod_{t=1}^{d} \mathbb{I}\Big[\,\sigma^{(t)}\Big(m_{c,i}^{(t)} - \theta^{(t)}\Big) \geq 0\Big] \;\in\; \{0,1\},$$

which reproduces the engine-specific rules of the Methods in a single expression. The set of accepted ranks and the selected rank are

$$S(c;\boldsymbol{\theta}) = \{\, i \in \{1,\ldots,n\} \;:\; F(\mathbf{m}_{c,i};\boldsymbol{\theta}) = 1 \,\}, \qquad r_{\text{best}}(c;\boldsymbol{\theta}) = \min S(c;\boldsymbol{\theta}),$$

with $r_{\text{best}}$ undefined when $S(c;\boldsymbol{\theta}) = \emptyset$. Selecting the *first* accepted rank, rather than the best-scoring accepted rank, keeps the engine's own ordering as a tie-breaker among poses that are all deemed acceptable.

**From acceptance to a ranking.** Let $g : \mathbb{R}^d \to \mathbb{R}$ be the scoring configuration under evaluation, i.e. the quantity by which compounds are finally ordered. We consider three configurations,

$$g_{\text{Aff}}(\mathbf{m}) = -\,E(\mathbf{m}), \qquad g_{\text{CNNaff}}(\mathbf{m}) = K(\mathbf{m}), \qquad g_{\text{CNNxAff}}(\mathbf{m}) = P(\mathbf{m}) \cdot K(\mathbf{m}),$$

where $E$, $P$ and $K$ denote the empirical affinity, `CNNpose` and `CNNaffinity` components of $\mathbf{m}$. The sign in $g_{\text{Aff}}$ makes larger values better for all three. The accepted compound set is

$$\mathcal{C}_{\text{acc}}(\boldsymbol{\theta}) = \{c \in \mathcal{C} : S(c;\boldsymbol{\theta}) \neq \emptyset\}\,.$$

For compounds in this set, the score induced by the re-ranking rule is

$$s(c;\boldsymbol{\theta}) = g\big(\mathbf{m}_{c,\,r_{\text{best}}(c;\boldsymbol{\theta})}\big)\,, \qquad c \in \mathcal{C}_{\text{acc}}(\boldsymbol{\theta}), \tag{S1}$$

The deposited `ProDock` branch writes `screened_withscore.csv` rows only for compounds with `Satisfied_count`>0. It assigns no fallback score to compounds with no accepted rank, and the subsequent GNINA and DiffDock merge is an inner join. Equation S1 therefore describes a ranking over retained compounds rather than over the complete library. The separate optimization and AUC script that produced the archived benchmark tables was not deposited, so its treatment of failed compounds cannot be independently verified. The retained set formulation is the most conservative reconstruction supported by the available implementation.

For any compound included in the baseline evaluation, the rank 1 score is

$$s_{\text{base}}(c) = g(\mathbf{m}_{c,1}) \,,$$

but the available files do not establish that the baseline and optimized rankings contain identical compound sets. The reported differences should therefore be interpreted as comparisons of archived pipeline outputs rather than as a verified paired comparison over every compound.

#### S1.2 Threshold optimization

Given a labeled training set $\mathcal{D}_{\text{train}} = \{(c, y_c)\}$ with $y_c \in \{0, 1\}$, the re-ranking rule is fitted by choosing the thresholds that maximize a screening objective $J$ evaluated on the induced ranking,

$$\boldsymbol{\theta}^* = \arg\max_{\boldsymbol{\theta}\in\Theta} \; J\Big(\{(s(c;\boldsymbol{\theta}),\, y_c)\}_{(c,y_c)\in\mathcal{D}_{\text{train}},\; c\in\mathcal{C}_{\text{acc}}(\boldsymbol{\theta})}\Big) \,.$$

Because $s(\cdot;\boldsymbol{\theta})$ depends on $\boldsymbol{\theta}$ only through the indicator functions in $F$, the objective is piecewise constant in $\boldsymbol{\theta}$ and has zero gradient almost everywhere. Gradient-based optimization is therefore inapplicable, which motivates the use of a sequential model-based (tree-structured Parzen estimator) sampler, as implemented in `Optuna` (cited in the main text). The procedure is summarized in Algorithm S1.

**Algorithm S1** `Optuna`-based threshold optimization for rank-resolved re-ranking.

**Require:** Descriptor tensor $\{\mathbf{m}_{c,i}\}$, labels $\{y_c\}$, scoring configuration $g$, search space $\Theta$, objective $J$, budget $T$
**Ensure:** Optimized thresholds $\boldsymbol{\theta}^*$ and retained held-out ranking

split $\mathcal{C}$ into $\mathcal{D}_{\text{train}}, \mathcal{D}_{\text{test}}$, stratified by $y$
**for** $\tau = 1, \ldots, T$ **do**
    $\boldsymbol{\theta}_\tau \leftarrow \text{Sample}(\Theta \mid \text{trial history})$ ▷ TPE sampler
    $\mathcal{R}_\tau \leftarrow \emptyset$
    **for all** $c \in \mathcal{D}_{\text{train}}$ **do**
        $S(c) \leftarrow \{\, i : F(\mathbf{m}_{c,i}; \boldsymbol{\theta}_\tau) = 1 \,\}$
        **if** $S(c) \neq \emptyset$ **then**
            add $(g(\mathbf{m}_{c,\min S(c)}), y_c)$ to the retained ranking $\mathcal{R}_\tau$
        **end if**
    **end for**
    $J_\tau \leftarrow J(\mathcal{R}_\tau)$
**end for**
$\boldsymbol{\theta}^* \leftarrow \boldsymbol{\theta}_{\tau^*}, \quad \tau^* = \arg\max_\tau J_\tau$
apply $\boldsymbol{\theta}^*$ *once* to the retained subset of $\mathcal{D}_{\text{test}}$

#### S1.3 Evaluation metrics

LogAUC is a logarithmically weighted ROC summary that emphasizes the low false positive rate region. A generic form is

$$\text{LogAUC}_\lambda \propto \int_\lambda^1 \text{TPR}(f) \frac{\mathrm{d}f}{f},$$

where the lower cutoff $\lambda$ and normalization are implementation choices. The generating optimization and metric code was not present in the deposited branch, and the archived tables do not encode these settings. The exact cutoff and normalization therefore cannot be recovered from the available materials. We report the archived LogAUC values without assigning an unverified numerical convention and compare only values carrying the same archived metric label.

For the separation coefficient of a single descriptor (main text), two properties of the estimator matter for its interpretation. It is a density-overlap rather than a rank-based statistic, so unlike ROC-AUC it is sensitive to the binning and to the range over which the histograms are formed: a single extreme value widens the common range, concentrates the remaining mass into few bins, and biases the coefficient toward zero. Descriptor columns must therefore be screened for sentinel or placeholder values before the coefficient is computed. It is also symmetric and unsigned, reporting the magnitude of the class difference without indicating which class scores higher.

#### S2 Optional ADMET reporting

`ProDock` includes an optional reporting module that submits retained ligands to the `ADMETLab 3.0` web server cited in the main text. The input is a `.csv` file containing compound names and `SMILES` strings, with an optional cluster label. For each compound, the workflow creates a ligand specific output directory, submits the `SMILES` string, and downloads the generated `.pdf` and `.csv` reports when available. Existing reports are detected before submission so interrupted or large jobs can resume without repeating successful predictions.

The module also generates a consolidated spreadsheet containing ligand names, `SMILES` strings, two dimensional structures, simplified radar plots for physicochemical descriptors, optional `ProLIF` interaction diagrams for the best satisfied `GNINA` pose, and the downloaded predictions. A plain `.csv` counterpart is exported for subsequent filtering or integration with external analysis tools. This optional module was not used in the `DUDE-Z` benchmark, and no ADMET results are reported in this study.

#### S3 Software and `Optuna` study configuration

All docking, descriptor calculation, optimization, and analysis used the software listed in Table S1, with random seeds fixed (conformer embedding and `GNINA` local docking both used seed 42) so that pose generation is reproducible from the deposited inputs.

Threshold optimization used the `Optuna` study configuration in Table S2. Each study maximized the training-split objective (ROC-AUC, PR-AUC, or LogAUC) with the de-

Table S1: Software used in the `ProDock` workflow and in the analyses reported here.

| Component | Software | Role |
|---|---|---|
| Global docking | `DiffDock` | Diffusion-based pose generation (10 poses/ligand) |
| Local docking | `GNINA` | CNN-scored local docking (seed 42, exhaustiveness 32, 10 poses) |
| Ligand preparation | `RDKit` | Protonation, 3D embedding (seed 42), MMFF minimization |
| Receptor preparation | `PyMOL`, `OpenMM` | Chain/cofactor extraction, protonation, AMBER14 minimization |
| Interactions | `ProLIF`, `MDAnalysis` | Protein–ligand interaction fingerprints |
| Electrostatics | `APBS`, `Open Babel` | Poisson–Boltzmann solvation energy |
| Optimization | `Optuna` | Threshold search (TPE sampler) |
| ADMET | `ADMETLab 3.0` | Optional property reporting |
| Visualization | `PyMOL`, `ChimeraX` | Pose overlays and interaction maps |

fault Tree-structured Parzen Estimator (TPE) sampler and no pruner, because each trial yields a single terminal objective value. Every threshold was a continuous variable (`trial.suggest_float`) sampled uniformly between the empirical minimum and maximum of that descriptor over the ten-pose pool of each molecule. Up to eight thresholds were optimized per study, four from `GNINA` and four from `DiffDock`; when the composite `CNNpose`×`CNNaffinity` score was used for ranking, `CNNpose` was removed from the threshold pool to avoid using the same quantity as both a ranking score and a filter. Critically, each of the nine scoring×metric configurations was optimized *independently for every target*, and the resulting threshold vector was frozen and applied once to that target's held-out test split.

Table S2: `Optuna` study configuration used for threshold optimization.

| Parameter | Value |
|---|---|
| Sampler | TPE (`Optuna` default) |
| Optimization direction | maximize |
| Pruner | none |
| Trials per study | 400 |
| Trial-level parallelism | 28 |
| Targets optimized in parallel | 28 |
| Per-target wall-clock timeout | 72 000 s (20 h) |
| Threshold variables | up to 8 (continuous, uniform) |
| Bounds per variable | empirical [min, max] of the ten-pose descriptor pool |
| Objective | ROC-AUC / PR-AUC / LogAUC (training split) |
| Scoring configuration | affinity / cnnaffinity / cnn-combined |

### S4 Benchmark Composition

Table S3 lists the make-up of the 43 `DUDE-Z` targets used in this work and the stratified 80/20 training/test partition used for threshold optimization. The benchmark contains 2,376 actives among 136,759 docked compounds, with a median of 49 actives per target (range 23–95) and a median decoy-to-active ratio of 56:1. This pronounced class imbalance is the reason PR-AUC and early-recognition metrics are emphasized in the main text.

Table S3: Composition of the DUDE-Z benchmark and the stratified 80/20 training/test partition used for threshold optimization. For each target: total docked compounds, experimentally supported actives, decoys, and the training-split size, training-split actives, and test-split size.

| Target | Compounds | Actives | Decoys | Training($n$) | Train act. | Test($n$) |
|---|---|---|---|---|---|---|
| AA2AR | 4585 | 87 | 4498 | 3668 | 70 | 917 |
| ABL1 | 3193 | 49 | 3144 | 2554 | 39 | 639 |
| ACES | 5022 | 84 | 4938 | 4017 | 67 | 1005 |
| ADA | 3360 | 62 | 3298 | 2688 | 50 | 672 |
| ADRB2 | 3303 | 55 | 3248 | 2642 | 44 | 661 |
| AMPC | 3193 | 48 | 3145 | 2554 | 38 | 639 |
| ANDR | 5038 | 94 | 4944 | 4030 | 75 | 1008 |
| CSF1R | 3553 | 56 | 3497 | 2842 | 45 | 711 |
| CXCR4 | 2229 | 30 | 2199 | 1783 | 24 | 446 |
| DEF | 2747 | 53 | 2694 | 2197 | 42 | 550 |
| DRD4 | 1873 | 24 | 1849 | 1498 | 19 | 375 |
| EGFR | 4315 | 68 | 4247 | 3452 | 54 | 863 |
| FA10 | 1320 | 23 | 1297 | 1056 | 18 | 264 |
| FA7 | 2180 | 32 | 2148 | 1744 | 26 | 436 |
| FABP4 | 2292 | 44 | 2248 | 1833 | 35 | 459 |
| FGFR1 | 3592 | 45 | 3547 | 2873 | 36 | 719 |
| FKB1A | 2305 | 46 | 2259 | 1844 | 37 | 461 |
| GLCM | 2185 | 37 | 2148 | 1748 | 30 | 437 |
| HDAC8 | 4173 | 75 | 4098 | 3338 | 60 | 835 |
| HIVPR | 3245 | 46 | 3199 | 2596 | 37 | 649 |
| HMDH | 1937 | 38 | 1899 | 1549 | 30 | 388 |
| HS90A | 3549 | 50 | 3499 | 2839 | 40 | 710 |
| ITAL | 2213 | 40 | 2173 | 1770 | 32 | 443 |
| KIT | 2644 | 46 | 2598 | 2115 | 37 | 529 |
| KITH | 3016 | 57 | 2959 | 2412 | 46 | 604 |
| LCK | 4013 | 66 | 3947 | 3210 | 53 | 803 |
| MAPK2 | 3972 | 75 | 3897 | 3177 | 60 | 795 |
| MK01 | 2291 | 43 | 2248 | 1832 | 34 | 459 |
| MT1 | 2176 | 28 | 2148 | 1740 | 22 | 436 |
| NRAM | 5621 | 91 | 5530 | 4496 | 73 | 1125 |
| PARP1 | 6210 | 95 | 6115 | 4968 | 76 | 1242 |
| PLK1 | 2171 | 42 | 2129 | 1736 | 34 | 435 |
| PPARA | 2228 | 48 | 2180 | 1782 | 38 | 446 |
| PTN1 | 3662 | 81 | 3581 | 2929 | 65 | 733 |
| PUR2 | 1175 | 40 | 1135 | 940 | 32 | 235 |
| RENI | 3819 | 85 | 3734 | 3055 | 68 | 764 |
| ROCK1 | 2000 | 42 | 1958 | 1600 | 34 | 400 |
| SRC | 4110 | 65 | 4045 | 3288 | 52 | 822 |
| THRB | 2690 | 44 | 2646 | 2152 | 35 | 538 |
| TRY1 | 3204 | 57 | 3147 | 2563 | 46 | 641 |
| TRYB1 | 1935 | 38 | 1897 | 1548 | 30 | 387 |
| UROK | 4568 | 73 | 4495 | 3654 | 58 | 914 |
| XIAP | 3852 | 74 | 3778 | 3081 | 59 | 771 |
| **Total** | **136759** | **2376** | **134383** | | | |

### S5 Score Distributions across All Pose Ranks and DUDE-Z Targets

Figures S1–S8 show the full distribution of every ProDock descriptor, separately for active and decoy compounds, across all ten pose ranks and all 43 DUDE-Z benchmark targets. Each panel row corresponds to one target and each column to one pose rank. The rank-averaged separation coefficients summarizing these distributions are reported in the main text.

Figures S1–S5 cover the GNINA scores, and Figures S6–S8 the DiffDock localization descriptors.

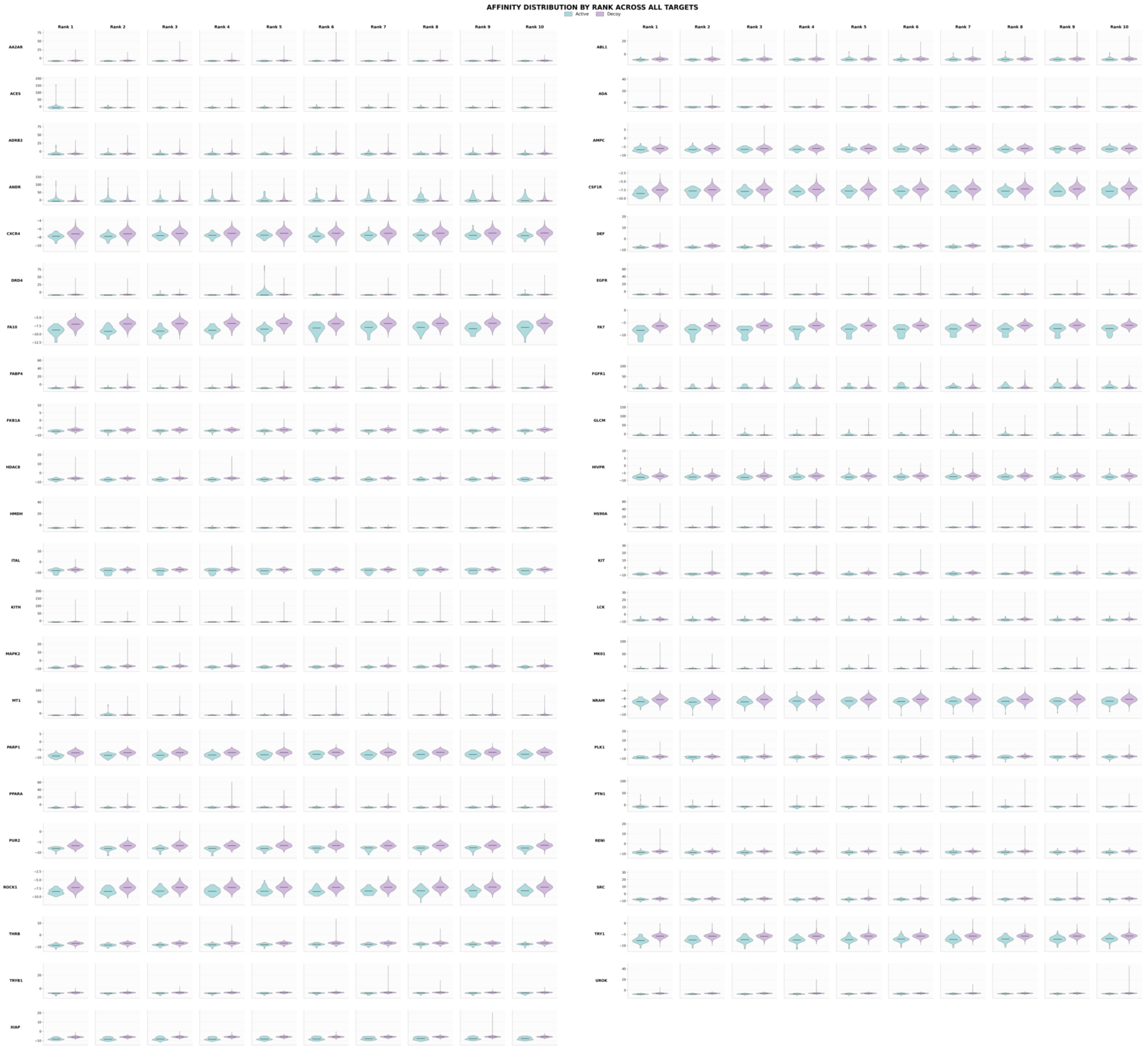


Figure S1: `GNINA Affinity` score distribution between actives (teal) and decoys (purple) across all ranks and targets.

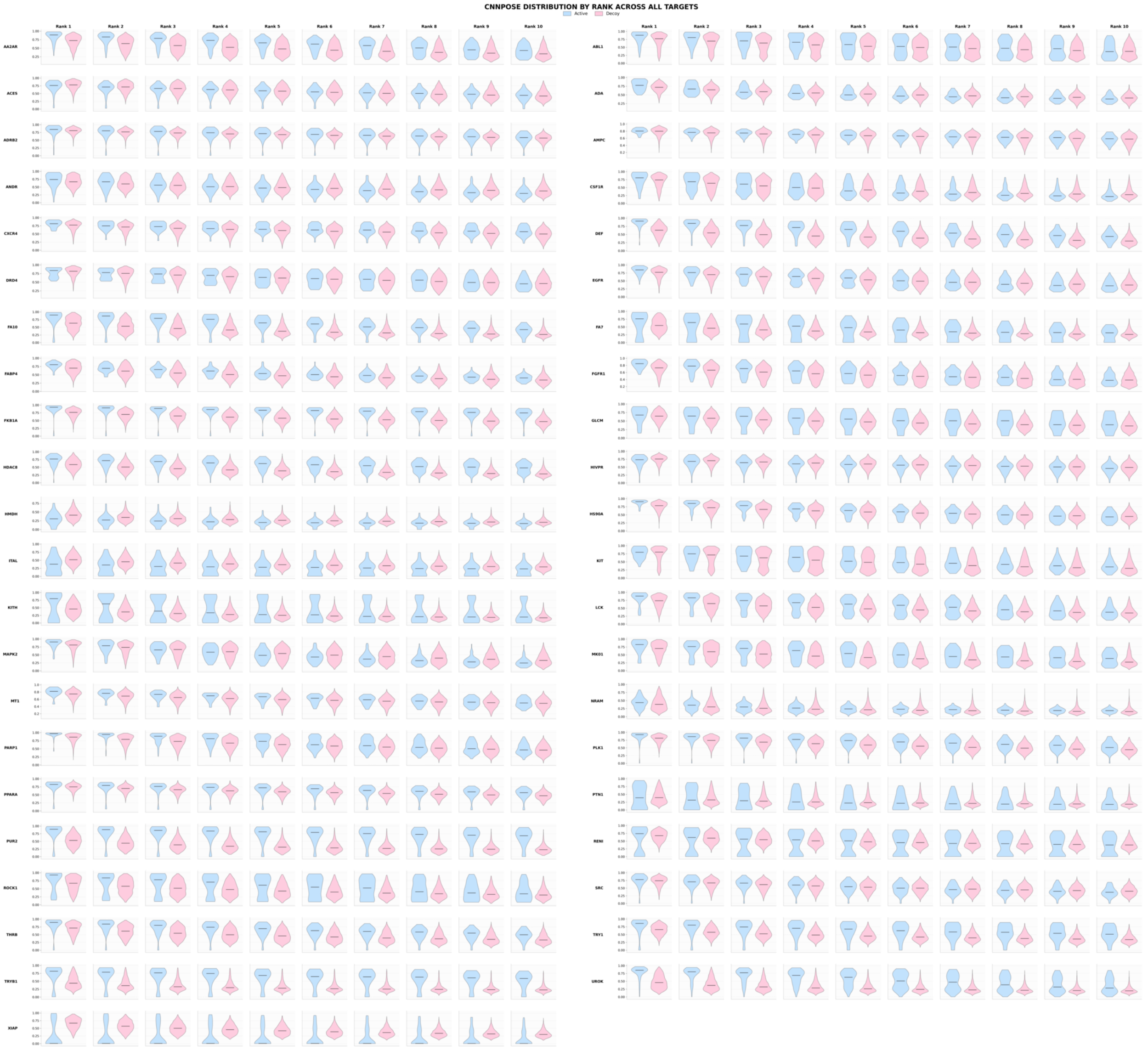


Figure S2: GNINA CNNpose score distribution between actives (blue) and decoys (pink) across all ranks and targets.

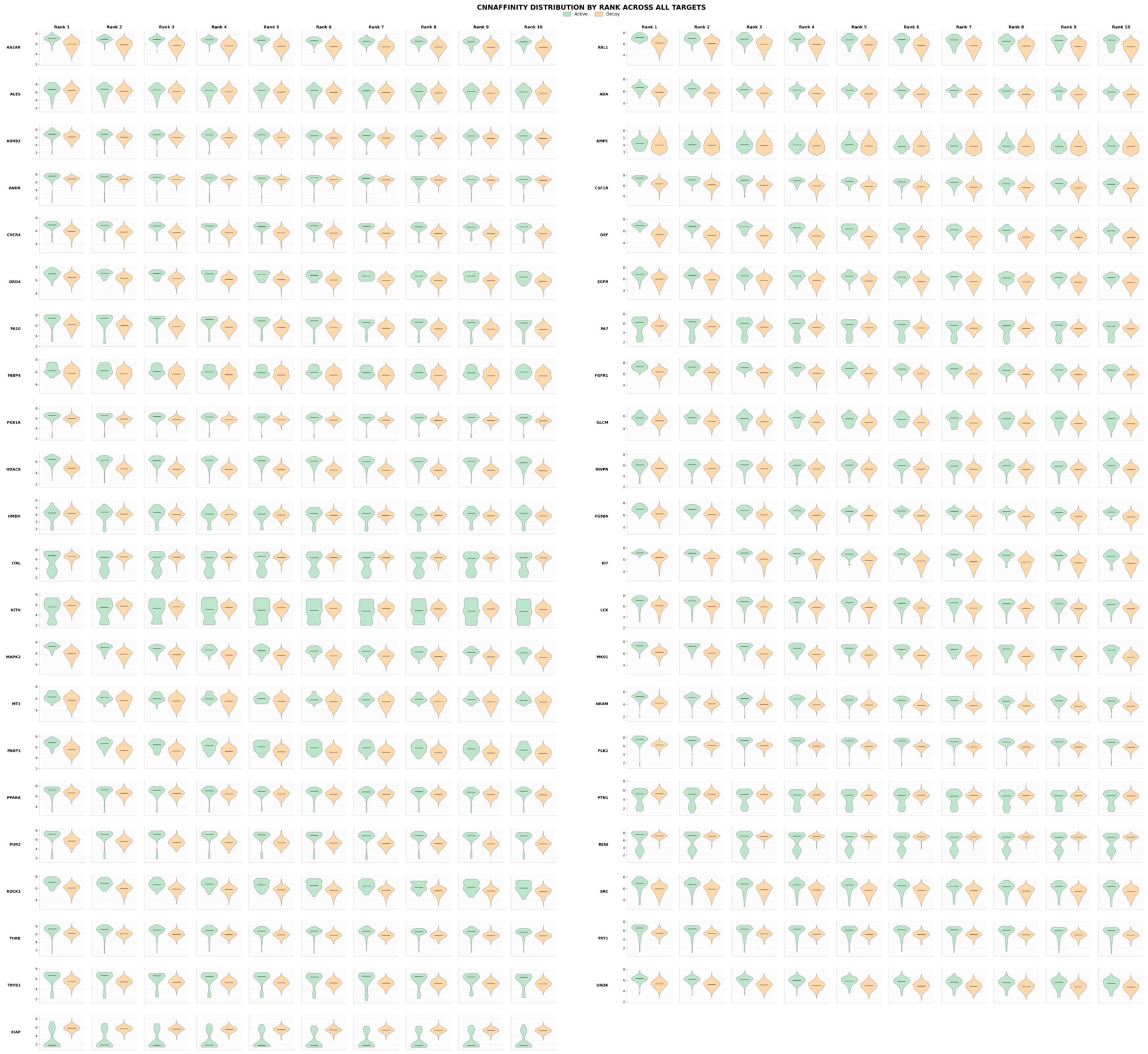


Figure S3: `GNINA CNNaffinity` score distribution between actives (green) and decoys (orange) across all ranks and targets.

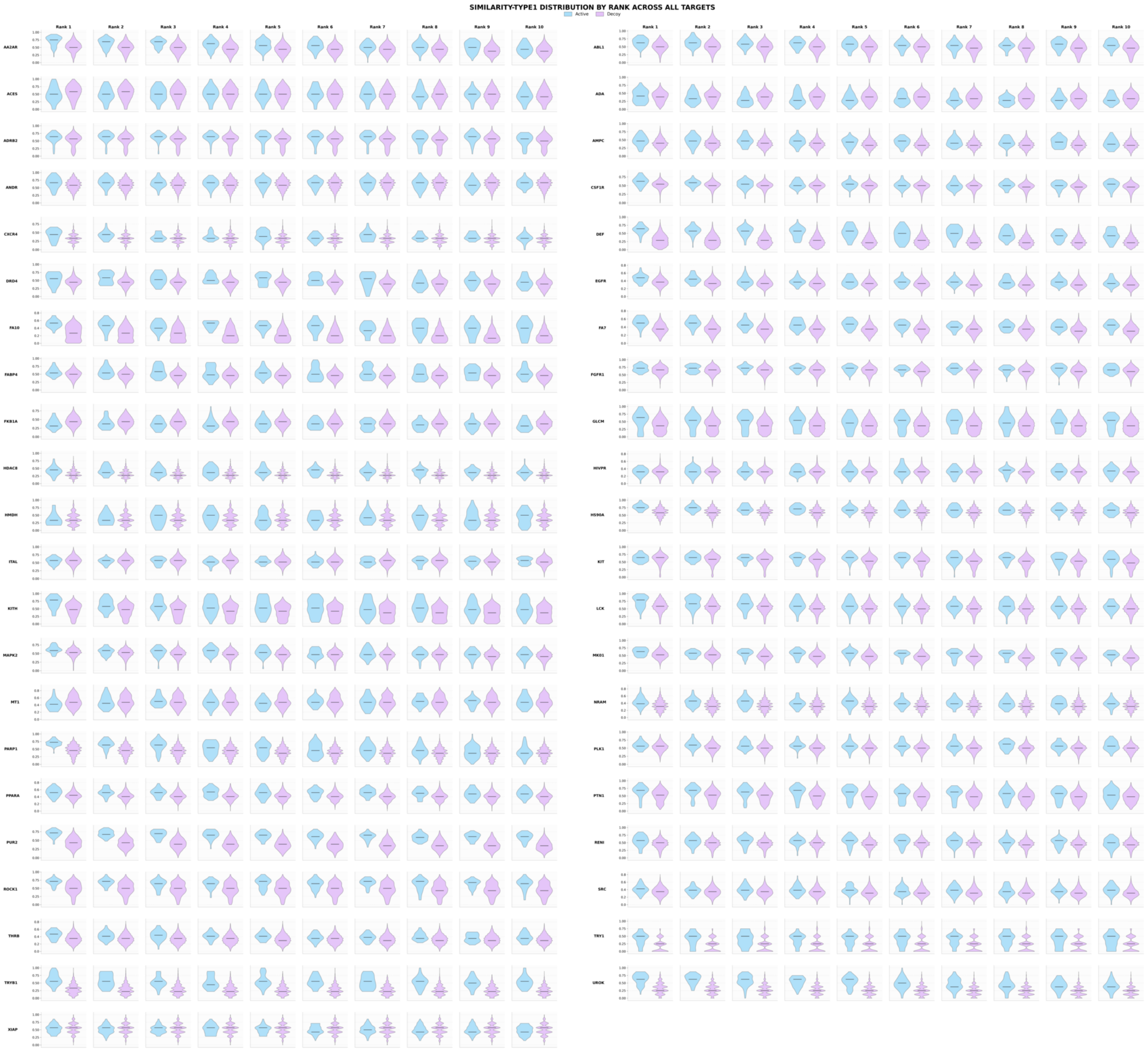


Figure S4: `GNINA Similarity type 1` score distribution between actives (blue) and decoys (purple) across all ranks and targets.

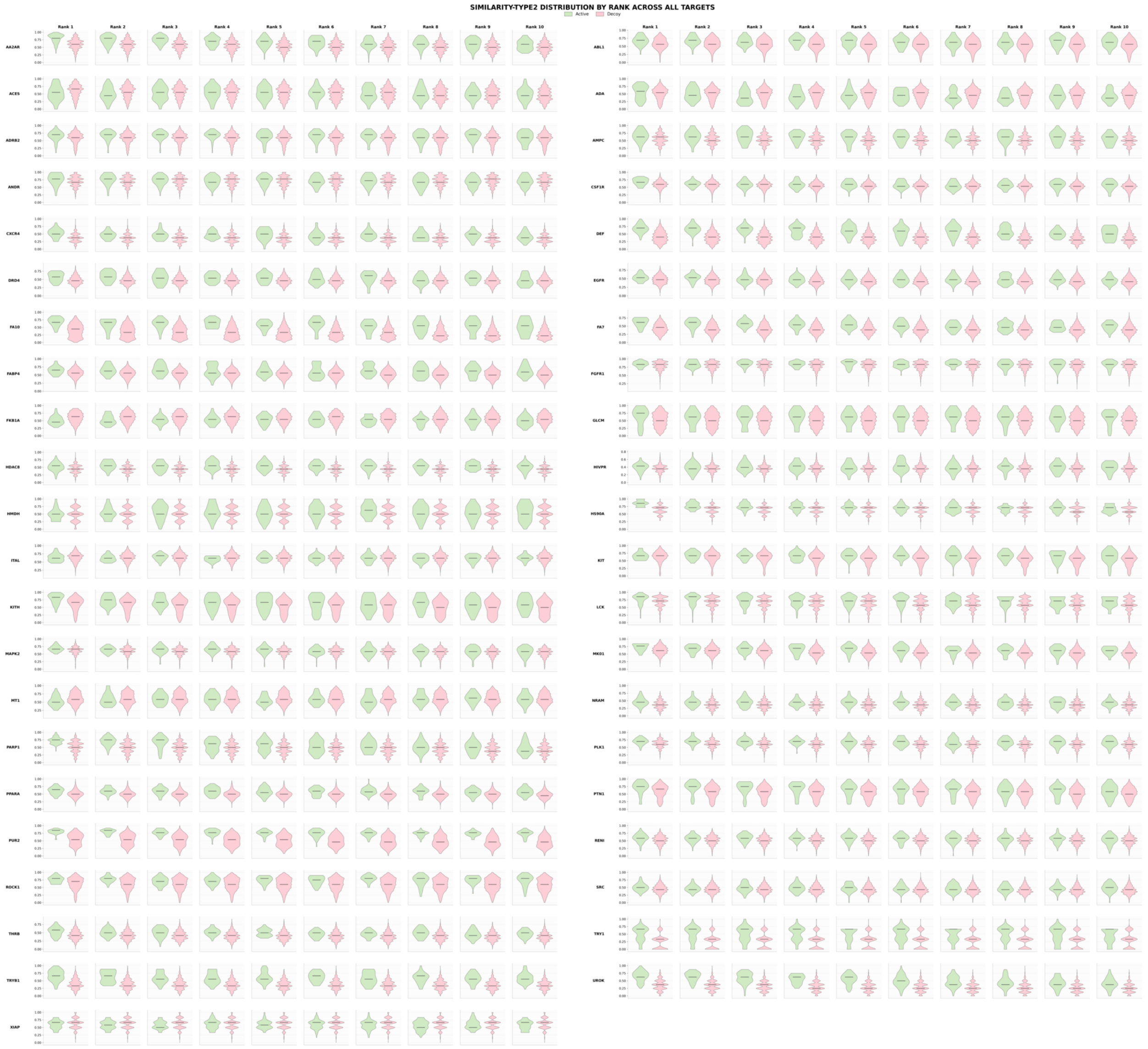

Figure S5: `GNINA Similarity type 2` score distribution between actives (green) and decoys (pink) across all ranks and targets.

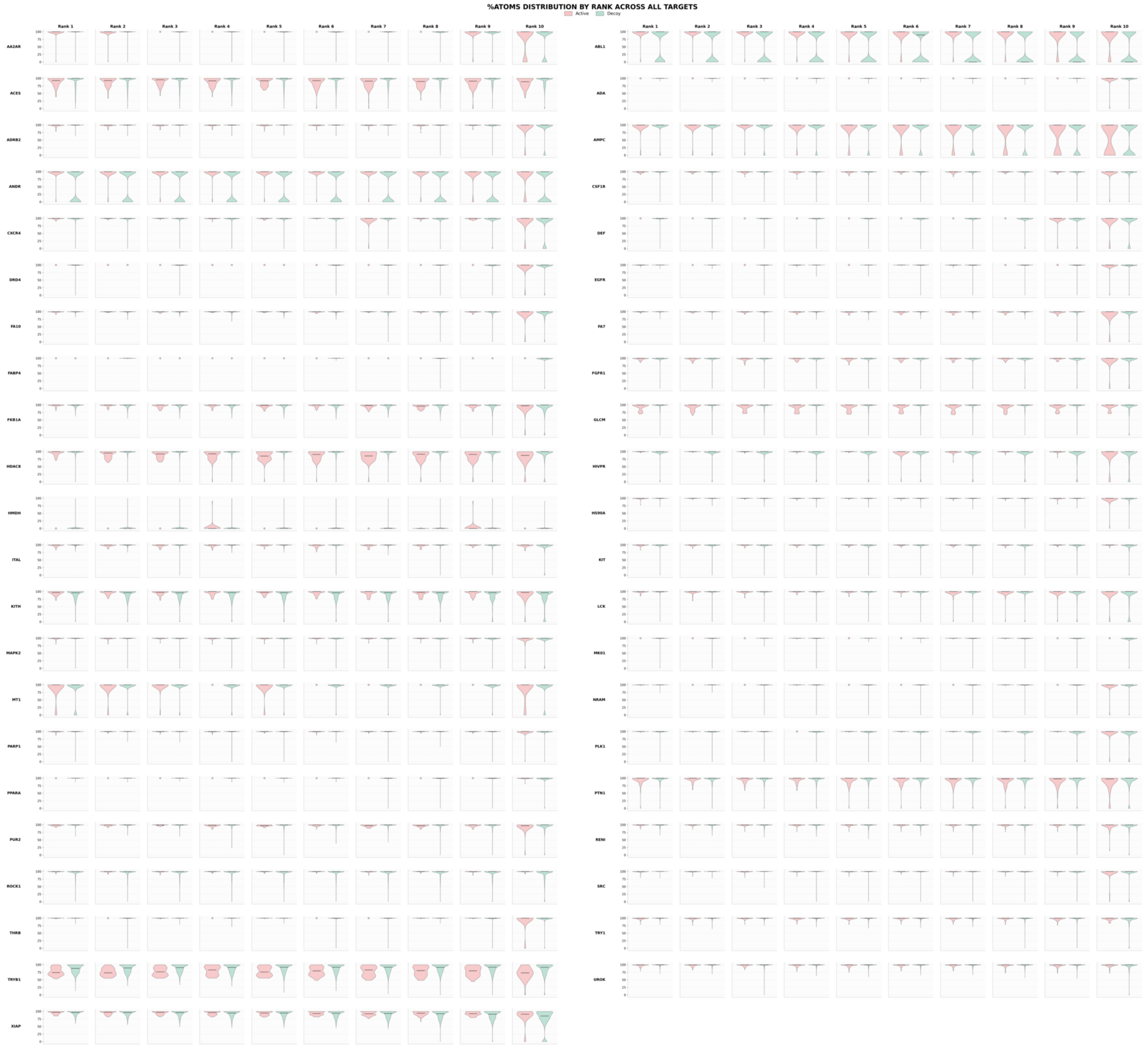


Figure S6: `DiffDock` %Atoms score distribution between actives (orange) and decoys (green) across all ranks and targets.

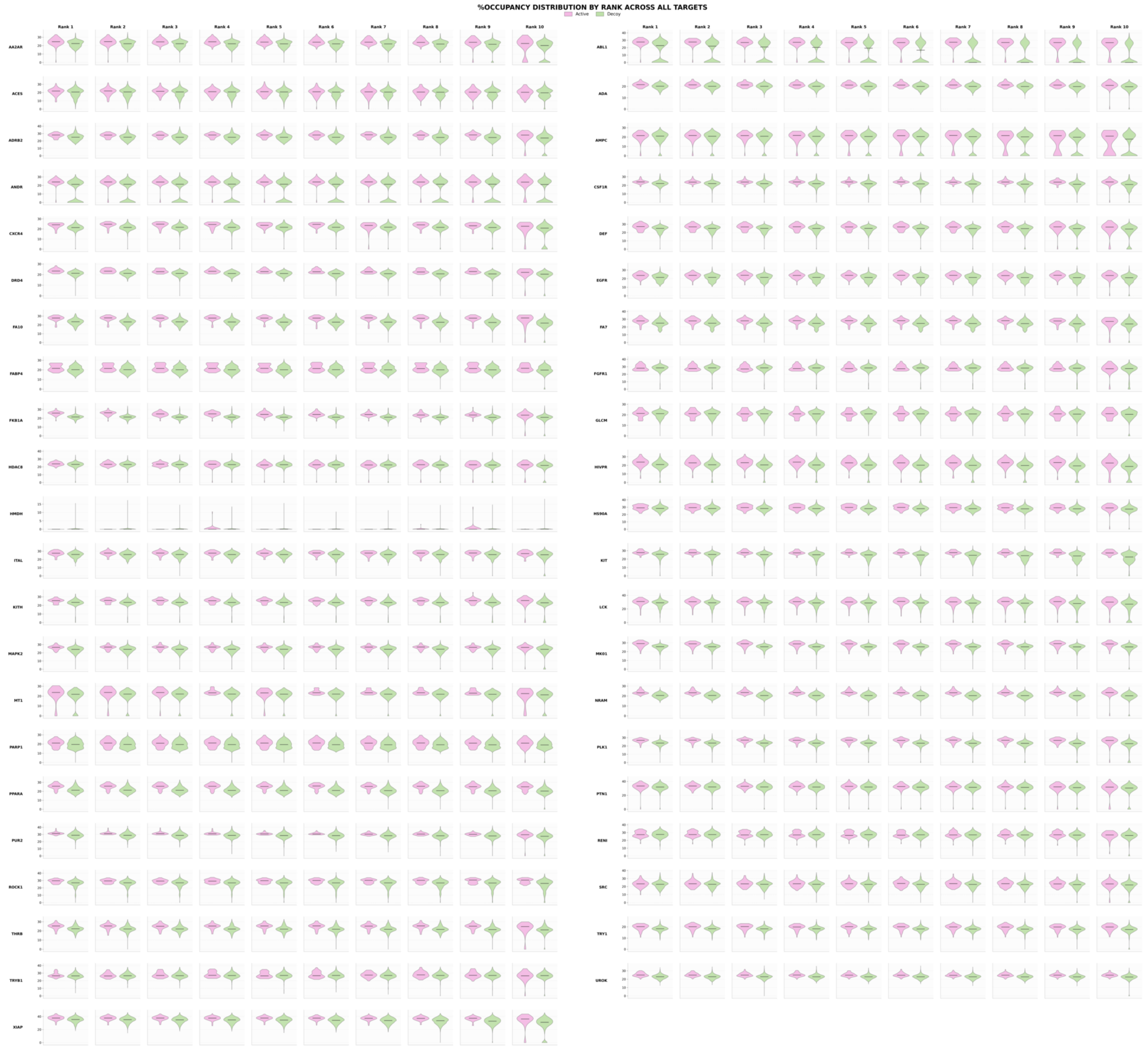


Figure S7: `DiffDock` normalized SASA localization score %Occupancy for actives (dark pink) and decoys (green) across all ranks and targets.

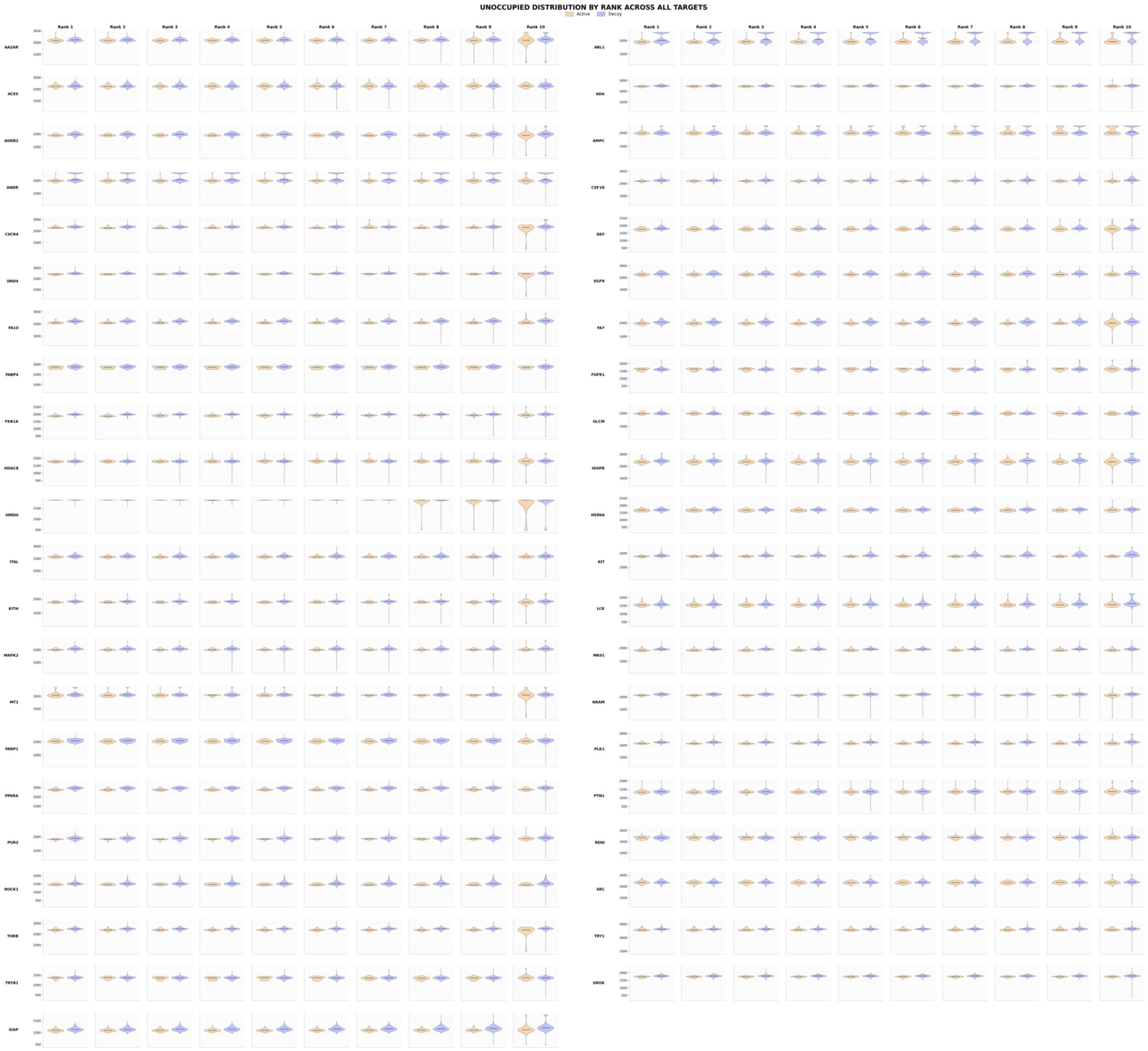


Figure S8: `DiffDock` residual SASA localization score `Unoccupied` for actives (yellow) and decoys (purple) across all ranks and targets. Raw `Unoccupied` and %Occupancy values are inversely related by construction. Their unsigned separation coefficients have Spearman $\rho = 0.99$ across target and rank combinations.

#### S6 Rank-Resolved Separation Coefficients

This section tabulates the two margins of the separation-coefficient data plotted in Figure 5 of the main text, so that the underlying per-target, per-rank values need not be consulted as raw data files. Table S4 gives the per-target values averaged over ranks (the values shown as the heatmap in Figure 5D), and Tables S5–S9 give the per-rank values averaged (mean and median) over targets, for every descriptor.

Table S4: Active-decoy separation coefficient for every `ProDock` descriptor and every target, averaged over the ten pose ranks, sorted by mean `GNINA` separation. These are the numerical values shown as the per-target heatmap in the descriptor-separation figure of the main text. Column keys: CNNaff, CNNpose, Aff = `GNINA` `CNNaffinity`, `CNNpose`, empirical affinity; Sim1, Sim2 = interaction-fingerprint similarity types 1 and 2; Conf = `DiffDock` confidence; %Occ, Unocc, %Atom = `DiffDock` occupancy, unoccupied area, and atom-localization. The electrostatic solvation energy is omitted (not computed in this run).

| Target | CNNaff | CNNpose | Aff | Sim1 | Sim2 | Conf | %Occ | Unocc | %Atom |
|---|---|---|---|---|---|---|---|---|---|
| PUR2 | 0.92 | 0.84 | 0.60 | 0.68 | 0.63 | 0.62 | 0.53 | 0.52 | 0.22 |
| XIAP | 0.68 | 0.77 | 0.61 | 0.12 | 0.13 | 0.69 | 0.39 | 0.39 | 0.25 |
| TRYB1 | 0.76 | 0.82 | 0.48 | 0.60 | 0.53 | 0.31 | 0.27 | 0.27 | 0.40 |
| FA10 | 0.77 | 0.66 | 0.60 | 0.54 | 0.53 | 0.51 | 0.73 | 0.71 | 0.15 |
| FA7 | 0.65 | 0.59 | 0.61 | 0.43 | 0.40 | 0.47 | 0.47 | 0.47 | 0.16 |
| HDAC8 | 0.73 | 0.50 | 0.55 | 0.33 | 0.30 | 0.29 | 0.16 | 0.12 | 0.45 |
| DEF | 0.69 | 0.54 | 0.53 | 0.56 | 0.49 | 0.64 | 0.31 | 0.31 | 0.03 |
| TRY1 | 0.63 | 0.53 | 0.56 | 0.36 | 0.37 | 0.32 | 0.45 | 0.44 | 0.16 |
| FKB1A | 0.51 | 0.71 | 0.45 | 0.25 | 0.25 | 0.66 | 0.64 | 0.65 | 0.24 |
| THRB | 0.62 | 0.51 | 0.54 | 0.31 | 0.31 | 0.49 | 0.59 | 0.59 | 0.03 |
| KITH | 0.55 | 0.64 | 0.47 | 0.37 | 0.26 | 0.45 | 0.36 | 0.36 | 0.21 |
| CXCR4 | 0.79 | 0.40 | 0.44 | 0.29 | 0.30 | 0.42 | 0.47 | 0.44 | 0.08 |
| ROCK1 | 0.53 | 0.55 | 0.49 | 0.42 | 0.34 | 0.48 | 0.39 | 0.38 | 0.13 |
| PLK1 | 0.78 | 0.43 | 0.32 | 0.29 | 0.27 | 0.41 | 0.56 | 0.56 | 0.04 |
| KIT | 0.67 | 0.35 | 0.48 | 0.28 | 0.26 | 0.35 | 0.49 | 0.48 | 0.08 |
| UROK | 0.48 | 0.54 | 0.46 | 0.57 | 0.52 | 0.52 | 0.44 | 0.44 | 0.14 |
| RENI | 0.47 | 0.56 | 0.44 | 0.26 | 0.20 | 0.56 | 0.27 | 0.26 | 0.09 |
| MK01 | 0.62 | 0.36 | 0.44 | 0.37 | 0.31 | 0.38 | 0.56 | 0.56 | 0.01 |
| ITAL | 0.52 | 0.50 | 0.40 | 0.23 | 0.18 | 0.36 | 0.34 | 0.34 | 0.13 |
| MAPK2 | 0.54 | 0.27 | 0.53 | 0.23 | 0.16 | 0.54 | 0.36 | 0.35 | 0.04 |
| PPARA | 0.46 | 0.46 | 0.41 | 0.38 | 0.38 | 0.49 | 0.59 | 0.59 | 0.01 |
| DRD4 | 0.53 | 0.43 | 0.35 | 0.37 | 0.30 | 0.36 | 0.50 | 0.50 | 0.00 |
| FABP4 | 0.37 | 0.36 | 0.54 | 0.35 | 0.30 | 0.38 | 0.37 | 0.36 | 0.00 |
| LCK | 0.53 | 0.32 | 0.42 | 0.24 | 0.21 | 0.36 | 0.29 | 0.29 | 0.08 |
| PARP1 | 0.43 | 0.36 | 0.43 | 0.34 | 0.27 | 0.54 | 0.23 | 0.23 | 0.06 |
| CSF1R | 0.58 | 0.31 | 0.33 | 0.24 | 0.20 | 0.36 | 0.45 | 0.45 | 0.09 |
| PTN1 | 0.42 | 0.49 | 0.28 | 0.32 | 0.28 | 0.27 | 0.19 | 0.16 | 0.15 |
| NRAM | 0.63 | 0.23 | 0.33 | 0.25 | 0.24 | 0.36 | 0.58 | 0.52 | 0.02 |
| AA2AR | 0.59 | 0.34 | 0.24 | 0.31 | 0.32 | 0.26 | 0.32 | 0.31 | 0.01 |
| FGFR1 | 0.56 | 0.32 | 0.28 | 0.28 | 0.20 | 0.44 | 0.31 | 0.30 | 0.25 |
| SRC | 0.52 | 0.25 | 0.38 | 0.27 | 0.22 | 0.41 | 0.21 | 0.21 | 0.05 |
| HMDH | 0.40 | 0.35 | 0.37 | 0.15 | 0.11 | 0.32 | 0.01 | 0.02 | 0.01 |
| GLCM | 0.45 | 0.46 | 0.21 | 0.30 | 0.26 | 0.32 | 0.40 | 0.39 | 0.26 |
| HS90A | 0.54 | 0.33 | 0.24 | 0.31 | 0.26 | 0.59 | 0.31 | 0.31 | 0.05 |
| ABL1 | 0.48 | 0.35 | 0.28 | 0.31 | 0.26 | 0.33 | 0.51 | 0.50 | 0.42 |
| HIVPR | 0.44 | 0.29 | 0.34 | 0.22 | 0.21 | 0.36 | 0.38 | 0.34 | 0.10 |
| MT1 | 0.45 | 0.36 | 0.21 | 0.25 | 0.22 | 0.37 | 0.40 | 0.40 | 0.03 |
| ADRB2 | 0.40 | 0.28 | 0.33 | 0.25 | 0.22 | 0.22 | 0.39 | 0.38 | 0.09 |
| EGFR | 0.45 | 0.27 | 0.24 | 0.25 | 0.21 | 0.34 | 0.31 | 0.31 | 0.01 |
| ADA | 0.43 | 0.28 | 0.21 | 0.22 | 0.23 | 0.38 | 0.33 | 0.32 | 0.01 |
| AMPC | 0.31 | 0.25 | 0.31 | 0.25 | 0.24 | 0.34 | 0.24 | 0.24 | 0.05 |
| ANDR | 0.34 | 0.23 | 0.17 | 0.13 | 0.10 | 0.38 | 0.46 | 0.45 | 0.36 |
| ACES | 0.24 | 0.24 | 0.15 | 0.16 | 0.15 | 0.19 | 0.19 | 0.17 | 0.30 |

The following tables give the per-rank values, averaged across targets.

Table S5: Mean and median active-decoy separation coefficients for the `GNINA Affinity` score across 43 targets at each pose rank.

| Pose rank | Mean separation | Median separation |
|---|---|---|
| 1 | 0.4319 | 0.4041 |
| 2 | 0.4295 | 0.4171 |
| 3 | 0.4138 | 0.4352 |
| 4 | 0.3934 | 0.3860 |
| 5 | 0.3986 | 0.3947 |
| 6 | 0.3830 | 0.3925 |
| 7 | 0.3870 | 0.3948 |
| 8 | 0.3693 | 0.3522 |
| 9 | 0.3828 | 0.3983 |
| 10 | 0.3778 | 0.3733 |

Table S6: Mean and median active-decoy separation coefficients for the `GNINA CNNpose` score across 43 targets at each pose rank.

| Pose rank | Mean separation | Median separation |
|---|---|---|
| 1 | 0.4712 | 0.4467 |
| 2 | 0.4677 | 0.4501 |
| 3 | 0.4529 | 0.4068 |
| 4 | 0.4468 | 0.4091 |
| 5 | 0.4371 | 0.3935 |
| 6 | 0.4302 | 0.4194 |
| 7 | 0.4204 | 0.3683 |
| 8 | 0.4116 | 0.3751 |
| 9 | 0.4047 | 0.3435 |
| 10 | 0.3967 | 0.3551 |

Table S7: Mean and median active-decoy separation coefficients for the `GNINA` `CNNaffinity` score across 43 targets at each pose rank.

| Pose rank | Mean separation | Median separation |
|---|---|---|
| 1 | 0.5782 | 0.5761 |
| 2 | 0.5730 | 0.5684 |
| 3 | 0.5588 | 0.5266 |
| 4 | 0.5558 | 0.5455 |
| 5 | 0.5404 | 0.5170 |
| 6 | 0.5420 | 0.5132 |
| 7 | 0.5327 | 0.5081 |
| 8 | 0.5244 | 0.5113 |
| 9 | 0.5263 | 0.5051 |
| 10 | 0.5181 | 0.4923 |

Table S8: Mean and median active-decoy separation coefficients for `Similarity type 1` and `Similarity type 2` across 43 targets at each pose rank.

| | Similarity type 1 | | Similarity type 2 | |
|---|---|---|---|---|
| Pose rank | Mean | Median | Mean | Median |
| 1 | 0.3758 | 0.3640 | 0.3353 | 0.2955 |
| 2 | 0.3456 | 0.3297 | 0.3081 | 0.2838 |
| 3 | 0.3336 | 0.3064 | 0.2920 | 0.2624 |
| 4 | 0.3242 | 0.3074 | 0.2915 | 0.2704 |
| 5 | 0.3167 | 0.2899 | 0.2842 | 0.2578 |
| 6 | 0.3023 | 0.2724 | 0.2601 | 0.2276 |
| 7 | 0.3089 | 0.2794 | 0.2822 | 0.2633 |
| 8 | 0.2953 | 0.2813 | 0.2494 | 0.2394 |
| 9 | 0.2900 | 0.2740 | 0.2584 | 0.2435 |
| 10 | 0.2791 | 0.2547 | 0.2609 | 0.2434 |

Table S9: Mean and median active-decoy separation coefficients for the `DiffDock` localization descriptors across 43 targets at each pose rank.

| | %Atoms | | %Occupancy | | **Unoccupied** | |
|---|---|---|---|---|---|---|
| **Pose rank** | **Mean** | **Median** | **Mean** | **Median** | **Mean** | **Median** |
| 1 | 0.1276 | 0.0742 | 0.3891 | 0.3898 | 0.3891 | 0.3898 |
| 2 | 0.1297 | 0.0873 | 0.3940 | 0.3687 | 0.3940 | 0.3687 |
| 3 | 0.1274 | 0.0814 | 0.3799 | 0.3599 | 0.3757 | 0.3523 |
| 4 | 0.1350 | 0.0886 | 0.3976 | 0.3741 | 0.3946 | 0.3695 |
| 5 | 0.1293 | 0.0875 | 0.4011 | 0.4045 | 0.3956 | 0.4045 |
| 6 | 0.1233 | 0.0755 | 0.3940 | 0.3818 | 0.3895 | 0.3818 |
| 7 | 0.1407 | 0.0703 | 0.3882 | 0.3925 | 0.3810 | 0.3917 |
| 8 | 0.1178 | 0.0530 | 0.3858 | 0.3914 | 0.3788 | 0.3806 |
| 9 | 0.1161 | 0.0625 | 0.3962 | 0.3984 | 0.3789 | 0.3529 |
| 10 | 0.1218 | 0.0817 | 0.3735 | 0.3782 | 0.3353 | 0.3482 |

#### S7 Per-Target Optimization Results

Tables S10–S12 give the held-out ROC-AUC, PR-AUC, and LogAUC for every target, before (rank-1 baseline) and after `Optuna`-optimized re-ranking, for the three scoring configurations; these are the per-target values summarized in the main-text results figures. Table S13 lists the optimized threshold vector selected on the training split for the `CNNaffinity`/PR-AUC configuration. Because thresholds were fitted independently per target, each descriptor spans a distribution of selected values rather than a single value; the %Atoms criterion, for example, was driven to its maximum of 100% in the majority of targets, indicating that it acts as a near-saturating pose-validity gate rather than a graded filter.

Table S10: Per-target held-out ROC-AUC before (rank-1 baseline) and after `Optuna`-optimized re-ranking, for the three scoring configurations. Bold marks a higher optimized value than the baseline.

| | Affinity | | `CNNaffinity` | | `CNNxAffinity` | |
|---|---|---|---|---|---|---|
| Target | Base | Opt | Base | Opt | Base | Opt |
| AA2AR | 0.731 | **0.835** | 0.930 | 0.856 | 0.887 | **0.900** |
| ABL1 | 0.687 | **0.723** | 0.873 | 0.859 | 0.710 | **0.790** |
| ACES | 0.579 | 0.548 | 0.620 | 0.504 | 0.615 | 0.606 |
| ADA | 0.648 | **0.712** | 0.759 | 0.745 | 0.660 | **0.686** |
| ADRB2 | 0.615 | **0.717** | 0.682 | **0.695** | 0.774 | **0.807** |
| AMPC | 0.491 | **0.621** | 0.434 | **0.592** | 0.512 | **0.606** |
| ANDR | 0.638 | 0.595 | 0.740 | 0.726 | 0.616 | **0.734** |
| CSF1R | 0.788 | **0.815** | 0.943 | 0.867 | 0.720 | **0.797** |
| CXCR4 | 0.794 | **0.928** | 0.990 | 0.889 | 0.890 | **0.950** |
| DEF | 0.869 | **0.886** | 0.955 | 0.881 | 0.923 | 0.882 |
| DRD4 | 0.881 | 0.787 | 0.777 | **0.809** | 0.690 | **0.715** |
| EGFR | 0.617 | **0.687** | 0.809 | 0.793 | 0.787 | 0.770 |
| FA10 | 0.962 | 0.885 | 0.744 | **0.786** | 0.769 | **0.981** |
| FA7 | 0.818 | 0.490 | 0.690 | 0.521 | 0.634 | 0.606 |
| FABP4 | 0.863 | 0.806 | 0.648 | 0.644 | 0.840 | 0.733 |
| FGFR1 | 0.569 | 0.508 | 0.887 | 0.638 | 0.818 | 0.598 |
| FKB1A | 0.808 | **0.983** | 0.767 | **0.976** | 0.941 | **0.992** |
| GLCM | 0.604 | **0.737** | 0.749 | 0.748 | 0.772 | 0.657 |
| HDAC8 | 0.843 | 0.759 | 0.920 | 0.766 | 0.860 | 0.701 |
| HIVPR | 0.622 | **0.648** | 0.597 | **0.703** | 0.505 | **0.634** |
| HMDH | 0.842 | 0.514 | 0.511 | **0.535** | 0.285 | **0.629** |
| HS90A | 0.650 | **0.813** | 0.813 | **0.904** | 0.917 | **0.947** |
| ITAL | 0.712 | 0.625 | 0.344 | **0.524** | 0.236 | **0.539** |
| KIT | 0.893 | **0.941** | 0.947 | 0.915 | 0.776 | **0.825** |
| KITH | 0.813 | 0.801 | 0.362 | **0.791** | 0.665 | **0.821** |
| LCK | 0.850 | 0.631 | 0.893 | 0.657 | 0.771 | 0.600 |
| MAPK2 | 0.869 | **0.884** | 0.896 | **0.909** | 0.840 | 0.832 |
| MK01 | 0.697 | **0.765** | 0.798 | **0.819** | 0.502 | **0.648** |
| MT1 | 0.627 | 0.603 | 0.508 | 0.498 | 0.650 | 0.555 |
| NRAM | 0.740 | **0.903** | 0.906 | **0.956** | 0.671 | **0.921** |
| PARP1 | 0.939 | **0.982** | 0.903 | **0.953** | 0.964 | **0.984** |
| PLK1 | 0.725 | 0.655 | 0.917 | **0.955** | 0.948 | 0.800 |
| PPARA | 0.863 | 0.847 | 0.775 | **0.872** | 0.832 | **0.870** |
| PTN1 | 0.739 | 0.555 | 0.464 | **0.534** | 0.569 | **0.599** |
| PUR2 | 0.817 | **0.987** | 0.974 | **0.993** | 0.989 | **0.995** |
| RENI | 0.754 | **0.832** | 0.625 | **0.774** | 0.631 | **0.776** |
| ROCK1 | 0.870 | 0.778 | 0.815 | **0.884** | 0.757 | **0.799** |
| SRC | 0.794 | **0.833** | 0.936 | 0.841 | 0.838 | 0.826 |
| THRB | 0.915 | 0.641 | 0.796 | 0.651 | 0.788 | 0.633 |
| TRY1 | 0.951 | 0.755 | 0.796 | **0.812** | 0.790 | 0.775 |
| TRYB1 | 0.830 | **0.980** | 0.971 | **0.987** | 0.986 | **0.994** |
| UROK | 0.915 | 0.890 | 0.865 | **0.882** | 0.900 | 0.881 |
| XIAP | 0.890 | 0.723 | 0.192 | **0.551** | 0.217 | **0.762** |
| Targets with a higher optimized value: Affinity 22/43, `CNNaffinity` 24/43, `CNNxAffinity` 27/43 | | | | | | |

Table S11: Per-target held-out PR-AUC before (rank-1 baseline) and after `Optuna`-optimized re-ranking. Bold marks a higher optimized value.

| | Affinity | | `CNNaffinity` | | `CNNxAffinity` | |
|---|---|---|---|---|---|---|
| Target | Base | Opt | Base | Opt | Base | Opt |
| AA2AR | 0.057 | **0.239** | 0.155 | **0.179** | 0.291 | 0.241 |
| ABL1 | 0.042 | 0.016 | 0.225 | **0.248** | 0.170 | 0.066 |
| ACES | 0.023 | 0.018 | 0.042 | 0.036 | 0.067 | 0.028 |
| ADA | 0.037 | **0.182** | 0.074 | **0.202** | 0.280 | **0.285** |
| ADRB2 | 0.071 | 0.017 | 0.030 | **0.048** | 0.055 | **0.129** |
| AMPC | 0.022 | **0.039** | 0.015 | **0.018** | 0.020 | **0.047** |
| ANDR | 0.211 | **0.294** | 0.170 | **0.269** | 0.150 | 0.133 |
| CSF1R | 0.065 | **0.162** | 0.322 | 0.245 | 0.349 | **0.354** |
| CXCR4 | 0.066 | **0.342** | 0.625 | 0.475 | 0.474 | **0.521** |
| DEF | 0.180 | **0.370** | 0.481 | 0.379 | 0.401 | **0.463** |
| DRD4 | 0.085 | **0.408** | 0.074 | **0.188** | 0.031 | **0.258** |
| EGFR | 0.025 | 0.016 | 0.238 | 0.208 | 0.220 | 0.170 |
| FA10 | 0.373 | **0.608** | 0.241 | **0.395** | 0.492 | **0.532** |
| FA7 | 0.276 | 0.095 | 0.044 | **0.067** | 0.044 | 0.039 |
| FABP4 | 0.151 | **0.302** | 0.044 | **0.237** | 0.063 | **0.237** |
| FGFR1 | 0.020 | 0.017 | 0.254 | 0.195 | 0.171 | 0.166 |
| FKB1A | 0.085 | **0.741** | 0.070 | **0.841** | 0.580 | **0.841** |
| GLCM | 0.185 | **0.338** | 0.110 | 0.016 | 0.223 | 0.195 |
| HDAC8 | 0.340 | 0.201 | 0.434 | 0.260 | 0.211 | 0.154 |
| HIVPR | 0.059 | 0.059 | 0.044 | 0.041 | 0.023 | **0.067** |
| HMDH | 0.129 | 0.036 | 0.030 | 0.021 | 0.015 | **0.021** |
| HS90A | 0.021 | **0.178** | 0.072 | **0.646** | 0.225 | **0.658** |
| ITAL | 0.278 | 0.018 | 0.024 | 0.018 | 0.013 | **0.018** |
| KIT | 0.331 | 0.043 | 0.329 | **0.569** | 0.161 | 0.126 |
| KITH | 0.314 | **0.417** | 0.019 | **0.107** | 0.233 | **0.603** |
| LCK | 0.091 | 0.025 | 0.324 | 0.219 | 0.345 | 0.106 |
| MAPK2 | 0.299 | **0.315** | 0.447 | **0.513** | 0.604 | 0.414 |
| MK01 | 0.070 | **0.265** | 0.227 | 0.220 | 0.026 | **0.097** |
| MT1 | 0.024 | 0.014 | 0.016 | 0.014 | 0.022 | 0.014 |
| NRAM | 0.052 | **0.289** | 0.137 | **0.455** | 0.035 | **0.289** |
| PARP1 | 0.400 | **0.570** | 0.360 | **0.416** | 0.634 | 0.436 |
| PLK1 | 0.088 | **0.141** | 0.294 | **0.546** | 0.592 | 0.556 |
| PPARA | 0.302 | **0.402** | 0.095 | **0.372** | 0.169 | **0.484** |
| PTN1 | 0.197 | 0.147 | 0.021 | **0.190** | 0.081 | 0.074 |
| PUR2 | 0.105 | **0.771** | 0.493 | **0.938** | 0.808 | **0.861** |
| RENI | 0.099 | **0.241** | 0.050 | **0.252** | 0.094 | **0.252** |
| ROCK1 | 0.181 | **0.274** | 0.402 | **0.632** | 0.658 | 0.559 |
| SRC | 0.140 | 0.095 | 0.362 | 0.208 | 0.414 | 0.322 |
| THRB | 0.385 | 0.341 | 0.147 | **0.280** | 0.201 | **0.222** |
| TRY1 | 0.460 | 0.350 | 0.229 | **0.405** | 0.485 | 0.426 |
| TRYB1 | 0.120 | **0.253** | 0.533 | 0.494 | 0.625 | **0.832** |
| UROK | 0.200 | **0.541** | 0.168 | **0.444** | 0.421 | **0.521** |
| XIAP | 0.695 | 0.085 | 0.013 | **0.128** | 0.026 | **0.142** |
| Targets with a higher optimized value: Affinity 25/43, `CNNaffinity` 27/43, `CNNxAffinity` 24/43 | | | | | | |

Table S12: Per-target held-out LogAUC before (rank-1 baseline) and after `Optuna`-optimized re-ranking. Bold marks a higher optimized value.

| | Affinity | | `CNNaffinity` | | `CNNxAffinity` | |
|---|---|---|---|---|---|---|
| Target | Base | Opt | Base | Opt | Base | Opt |
| AA2AR | 0.700 | **0.845** | 0.916 | 0.897 | 0.858 | **0.887** |
| ABL1 | 0.639 | **0.708** | 0.827 | 0.822 | 0.673 | **0.790** |
| ACES | 0.553 | 0.535 | 0.592 | 0.535 | 0.587 | 0.570 |
| ADA | 0.614 | 0.611 | 0.707 | 0.701 | 0.634 | **0.754** |
| ADRB2 | 0.575 | **0.603** | 0.639 | 0.619 | 0.732 | 0.711 |
| AMPC | 0.443 | **0.609** | 0.393 | **0.579** | 0.467 | **0.596** |
| ANDR | 0.604 | **0.694** | 0.714 | **0.813** | 0.591 | **0.698** |
| CSF1R | 0.745 | **0.753** | 0.898 | 0.818 | 0.676 | **0.811** |
| CXCR4 | 0.713 | **0.791** | 0.827 | **0.866** | 0.808 | **0.921** |
| DEF | 0.839 | **0.974** | 0.909 | **0.957** | 0.879 | **0.966** |
| DRD4 | 0.800 | **0.829** | 0.680 | **0.757** | 0.602 | **0.662** |
| EGFR | 0.585 | **0.677** | 0.780 | 0.776 | 0.753 | **0.758** |
| FA10 | 0.893 | 0.879 | 0.647 | **0.805** | 0.678 | **0.827** |
| FA7 | 0.720 | 0.648 | 0.617 | 0.576 | 0.554 | 0.478 |
| FABP4 | 0.810 | **0.867** | 0.594 | **0.674** | 0.785 | 0.693 |
| FGFR1 | 0.520 | 0.500 | 0.834 | 0.695 | 0.770 | 0.557 |
| FKB1A | 0.755 | **0.971** | 0.714 | **0.982** | 0.889 | **0.979** |
| GLCM | 0.521 | **0.633** | 0.679 | 0.678 | 0.721 | 0.492 |
| HDAC8 | 0.817 | 0.749 | 0.886 | 0.755 | 0.830 | 0.687 |
| HIVPR | 0.568 | **0.717** | 0.546 | **0.718** | 0.451 | **0.711** |
| HMDH | 0.782 | 0.500 | 0.453 | **0.595** | 0.242 | **0.479** |
| HS90A | 0.623 | **0.872** | 0.764 | **0.853** | 0.866 | **0.936** |
| ITAL | 0.651 | 0.559 | 0.295 | **0.482** | 0.185 | **0.491** |
| KIT | 0.841 | **0.893** | 0.895 | 0.873 | 0.724 | **0.811** |
| KITH | 0.768 | **0.803** | 0.319 | **0.785** | 0.619 | **0.767** |
| LCK | 0.815 | 0.612 | 0.857 | 0.583 | 0.733 | 0.625 |
| MAPK2 | 0.841 | **0.850** | 0.864 | **0.883** | 0.807 | **0.814** |
| MK01 | 0.644 | **0.719** | 0.765 | 0.677 | 0.451 | 0.436 |
| MT1 | 0.554 | **0.575** | 0.439 | **0.603** | 0.582 | 0.522 |
| NRAM | 0.716 | **0.894** | 0.879 | **0.934** | 0.642 | **0.913** |
| PARP1 | 0.914 | **0.974** | 0.877 | **0.965** | 0.938 | **0.974** |
| PLK1 | 0.666 | **0.703** | 0.856 | **0.906** | 0.886 | 0.777 |
| PPARA | 0.813 | **0.837** | 0.727 | **0.840** | 0.784 | **0.841** |
| PTN1 | 0.715 | 0.550 | 0.438 | **0.534** | 0.538 | **0.577** |
| PUR2 | 0.759 | **0.927** | 0.921 | **0.929** | 0.868 | **0.986** |
| RENI | 0.725 | **0.776** | 0.597 | **0.676** | 0.602 | 0.594 |
| ROCK1 | 0.838 | 0.706 | 0.752 | **0.875** | 0.696 | **0.731** |
| SRC | 0.765 | **0.811** | 0.898 | **0.910** | 0.802 | 0.774 |
| THRB | 0.863 | 0.638 | 0.743 | 0.633 | 0.733 | 0.638 |
| TRY1 | 0.906 | 0.778 | 0.789 | **0.800** | 0.746 | 0.726 |
| TRYB1 | 0.761 | **0.860** | 0.912 | **0.980** | 0.920 | **0.979** |
| UROK | 0.887 | **0.896** | 0.836 | **0.878** | 0.868 | 0.858 |
| XIAP | 0.867 | 0.549 | 0.163 | **0.613** | 0.184 | **0.565** |
| Targets with a higher optimized value: Affinity 29/43, `CNNaffinity` 28/43, `CNNxAffinity` 27/43 | | | | | | |

Table S13: `Optuna`-optimized threshold values selected on the training split for the `CNNaffinity`/PR-AUC configuration, one row per target. Thresholds for the other eight scoring×metric configurations are provided as `.csv` files in the data repository. Occ = occupied-area threshold; %Occ = percent occupancy; %Atom = percent of ligand atoms in the site.

| Target | CNNpose | Sim1 | Sim2 | Conf | Occ | %Occ | %Atom |
|---|---|---|---|---|---|---|---|
| AA2AR | 0.91 | 0.20 | 0.87 | -4.41 | 2327.21 | 14.78 | 100.00 |
| ABL1 | 0.14 | 0.68 | 0.63 | -4.45 | 2292.21 | 17.14 | 13.48 |
| ACES | 0.41 | 0.07 | 0.41 | -1.35 | 2697.18 | 18.83 | 61.39 |
| ADA | 0.83 | 0.31 | 0.55 | -0.24 | 2509.29 | 20.26 | 100.00 |
| ADRB2 | 0.66 | 0.54 | 0.15 | -0.52 | 2019.05 | 14.58 | 100.00 |
| AMPC | 0.78 | 0.04 | 0.80 | -1.96 | 2503.94 | 12.98 | 100.00 |
| ANDR | 0.33 | 0.21 | 0.45 | -2.56 | 2366.11 | 4.48 | 91.20 |
| CSF1R | 0.12 | 0.62 | 0.49 | -0.68 | 2234.90 | 20.02 | 100.00 |
| CXCR4 | 0.65 | 0.41 | 0.19 | -3.01 | 2409.70 | 19.05 | 100.00 |
| DEF | 0.55 | 0.24 | 0.07 | -0.14 | 1963.02 | 15.87 | 100.00 |
| DRD4 | 0.66 | 0.66 | 0.55 | -2.05 | 2431.31 | 16.44 | 100.00 |
| EGFR | 0.36 | 0.25 | 0.22 | -0.76 | 2431.08 | 19.29 | 100.00 |
| FA10 | 0.57 | 0.08 | 0.06 | -1.01 | 2115.04 | 12.92 | 100.00 |
| FA7 | 0.68 | 0.47 | 0.45 | -0.20 | 2346.64 | 24.35 | 100.00 |
| FABP4 | 0.40 | 0.82 | 0.41 | -1.49 | 3008.00 | 21.93 | 100.00 |
| FGFR1 | 0.69 | 0.74 | 0.81 | -2.97 | 1778.25 | 13.17 | 100.00 |
| FKB1A | 0.82 | 0.10 | 0.06 | -0.94 | 1895.10 | 22.47 | 92.82 |
| GLCM | 0.64 | 0.87 | 0.49 | -1.24 | 2183.05 | 19.96 | 100.00 |
| HDAC8 | 0.25 | 0.34 | 0.30 | -2.72 | 1969.61 | 22.79 | 100.00 |
| HIVPR | 0.06 | 0.03 | 0.22 | -0.49 | 2738.69 | 6.29 | 100.00 |
| HMDH | 0.64 | 0.05 | 0.49 | -0.87 | 1882.86 | 0.00 | 0.00 |
| HS90A | 0.22 | 0.74 | 0.78 | -0.29 | 1807.28 | 19.05 | 100.00 |
| ITAL | 0.54 | 0.30 | 0.54 | -1.05 | 2213.65 | 22.25 | 100.00 |
| KIT | 0.31 | 0.42 | 0.14 | -6.84 | 1842.34 | 25.72 | 100.00 |
| KITH | 0.92 | 0.62 | 0.32 | -3.64 | 1976.85 | 24.58 | 71.89 |
| LCK | 0.08 | 0.56 | 0.74 | -0.87 | 1851.64 | 19.39 | 100.00 |
| MAPK2 | 0.39 | 0.24 | 0.33 | -0.24 | 2229.22 | 15.71 | 100.00 |
| MK01 | 0.16 | 0.20 | 0.22 | -2.15 | 1905.74 | 19.02 | 100.00 |
| MT1 | 0.34 | 0.83 | 0.75 | -5.21 | 2119.89 | 27.19 | 100.00 |
| NRAM | 0.07 | 0.32 | 0.41 | -0.41 | 2470.19 | 20.12 | 100.00 |
| PARP1 | 0.85 | 0.59 | 0.56 | 0.09 | 2377.30 | 1.03 | 100.00 |
| PLK1 | 0.35 | 0.16 | 0.46 | -0.88 | 2237.11 | 20.75 | 100.00 |
| PPARA | 0.75 | 0.49 | 0.11 | -0.61 | 3143.51 | 18.05 | 100.00 |
| PTN1 | 0.82 | 0.15 | 0.05 | -0.25 | 1736.67 | 25.60 | 88.45 |
| PUR2 | 0.61 | 0.27 | 0.70 | -0.88 | 2330.43 | 28.88 | 100.00 |
| RENI | 0.81 | 0.65 | 0.43 | -1.17 | 2221.97 | 19.44 | 94.14 |
| ROCK1 | 0.75 | 0.58 | 0.39 | -5.54 | 1612.31 | 9.70 | 100.00 |
| SRC | 0.62 | 0.08 | 0.27 | -1.16 | 2940.82 | 8.81 | 100.00 |
| THRB | 0.57 | 0.24 | 0.08 | 0.03 | 2430.13 | 13.51 | 100.00 |
| TRY1 | 0.77 | 0.37 | 0.42 | -1.78 | 2803.55 | 15.21 | 100.00 |
| TRYB1 | 0.58 | 0.58 | 0.04 | -4.93 | 1513.69 | 15.78 | 81.65 |
| UROK | 0.06 | 0.68 | 0.56 | -0.55 | 1834.47 | 18.20 | 100.00 |
| XIAP | 0.55 | 0.04 | 0.29 | 0.30 | 1318.83 | 27.48 | 97.45 |

### S8 Structural Rescue Analysis

Table S14 reports the structural comparison underlying the main-text pose-recovery analysis: for the 18 rescued held-out actives of the three targets with an available co-crystal structure (NRAM, XIAP, HIVPR), it gives the native-site occupancy, pose–reference centroid distance, and interaction-fingerprint similarity of the rank-1 pose and of the pose selected by re-ranking.

Table S14: Structural comparison of the rank-1 and re-ranked (selected) `GNINA` poses against the co-crystal ligand, for the 18 rescued held-out actives of the three targets with an available co-crystal structure. Occupancy is the fraction of ligand heavy atoms within 2 Å of any co-crystal ligand atom; $d_{\mathrm{cent}}$ is the pose–reference centroid distance; $S_2$ is type-2 `ProLIF` similarity. Poses were compared in the original docking frame without superposition.

| | | Sel. | Occupancy | | $d_{\mathrm{cent}}$ (Å) | | $S_2$ | |
|---|---|---|---|---|---|---|---|---|
| Target | Compound | rank | r1 | sel | r1 | sel | r1 | sel |
| NRAM | CHEMBL167253 | 3 | 0.59 | 0.38 | 1.96 | 3.40 | 0.36 | 0.46 |
| NRAM | CHEMBL373336 | 5 | 0.68 | 0.86 | 2.18 | 1.37 | 0.36 | 0.46 |
| NRAM | CHEMBL311059 | 2 | 0.86 | 0.86 | 0.47 | 1.61 | 0.18 | 0.46 |
| NRAM | CHEMBL418456 | 2 | 0.53 | 0.60 | 3.01 | 1.71 | 0.64 | 0.82 |
| NRAM | CHEMBL190819 | 5 | 0.95 | 1.00 | 0.68 | 1.15 | 0.55 | 0.64 |
| NRAM | CHEMBL350298 | 9 | 0.61 | 0.77 | 2.30 | 2.15 | 0.27 | 0.55 |
| NRAM | CHEMBL254679 | 2 | 0.95 | 0.86 | 1.36 | 1.03 | 0.27 | 0.55 |
| NRAM | CHEMBL436250 | 10 | 0.83 | 0.87 | 1.46 | 1.14 | 0.46 | 0.55 |
| NRAM | CHEMBL164976 | 2 | 0.95 | 0.86 | 1.04 | 0.96 | 0.36 | 0.55 |
| NRAM | CHEMBL81777 | 3 | 0.62 | 0.95 | 3.62 | 1.10 | 0.36 | 0.46 |
| XIAP | CHEMBL375784 | 2 | 0.55 | 0.82 | 4.61 | 2.31 | 0.33 | 0.67 |
| XIAP | CHEMBL584397 | 3 | 0.61 | 0.88 | 4.43 | 3.75 | 0.67 | 0.83 |
| XIAP | CHEMBL573787 | 3 | 0.62 | 0.85 | 4.66 | 4.14 | 0.50 | 0.67 |
| XIAP | CHEMBL485721 | 2 | 0.75 | 0.97 | 5.35 | 4.18 | 0.33 | 0.50 |
| HIVPR | CHEMBL119970 | 2 | 0.56 | 0.72 | 2.37 | 0.99 | 0.50 | 0.79 |
| HIVPR | CHEMBL62390 | 3 | 0.55 | 0.79 | 4.07 | 1.56 | 0.43 | 0.50 |
| HIVPR | CHEMBL148371 | 2 | 0.79 | 0.83 | 3.25 | 2.46 | 0.36 | 0.43 |
| HIVPR | CHEMBL143758 | 3 | 0.50 | 0.83 | 3.91 | 2.53 | 0.29 | 0.36 |
